# ANKRD5: a key component of the axoneme required for sperm motility and male fertility

**DOI:** 10.1101/2024.12.03.626701

**Authors:** Shuntai Yu, Guoliang Yin, Peng Jin, Weilin Zhang, Yingchao Tian, Xiaotong Xu, Tianyu Shao, Yushan Li, Fei Sun, Yun Zhu, Fengchao Wang

**Affiliations:** Academy for Advanced Interdisciplinary Studies, Peking University, Beijing 100871, China; National Institute of Biological Sciences (NIBS), Beijing 102206, China; Tsinghua Institute of Multidisciplinary Biomedical Research, Tsinghua University, Beijing 102206, China; Key Laboratory of Biomacromolecules, CAS Center for Excellence in Biomacromolecules, Institute of Biophysics, Chinese Academy of Sciences, Beijing 100101, China; University of Chinese Academy of Sciences, Beijing 100049, China; The School of Public Health, Xinxiang Medical University, Xinxiang 453003, China

**Keywords:** sperm motility, ANKRD5, axoneme, N-DRC, male infertility

## Abstract

Sperm motility is essential for male fertility and depends on the structural integrity of the sperm axoneme, which features a canonical "9+2" microtubule arrangement. This structure comprises nine outer doublet microtubules (DMTs) that are associated with various macromolecular complexes. Among them, the nexin-dynein regulatory complex (N-DRC) forms crossbridges between adjacent DMTs, contributing to their stabilization and enabling flagellar bending. In this study, we investigated Ankyrin repeat domain 5 (ANKRD5, also known as ANK5 or ANKEF1), a protein highly expressed in the sperm axoneme. We found that ANKRD5 interacts with DRC5/TCTE1 and DRC4/GAS8, two key components of the N-DRC, and these interactions occur independently of calcium regulation. Male *Ankrd5*^-/-^ mice exhibited impaired sperm motility and infertility. Cryo-electron tomography revealed a typical "9+2" axoneme structure with intact DMTs in *Ankrd5* null sperm; however, the DMTs showed pronounced morphological variability and increased structural heterogeneity. Notably, ANKRD5 deficiency did not alter ATP levels, reactive oxygen species (ROS) levels, or mitochondrial membrane potential. These findings suggest that ANKRD5 may attenuate the N-DRC’s mechanical buffering-akin to a "car bumper"-between adjacent DMTs, thereby compromising axonemal stability under high mechanical stress during vigorous flagellar beating.

**Graphic Abstract:** 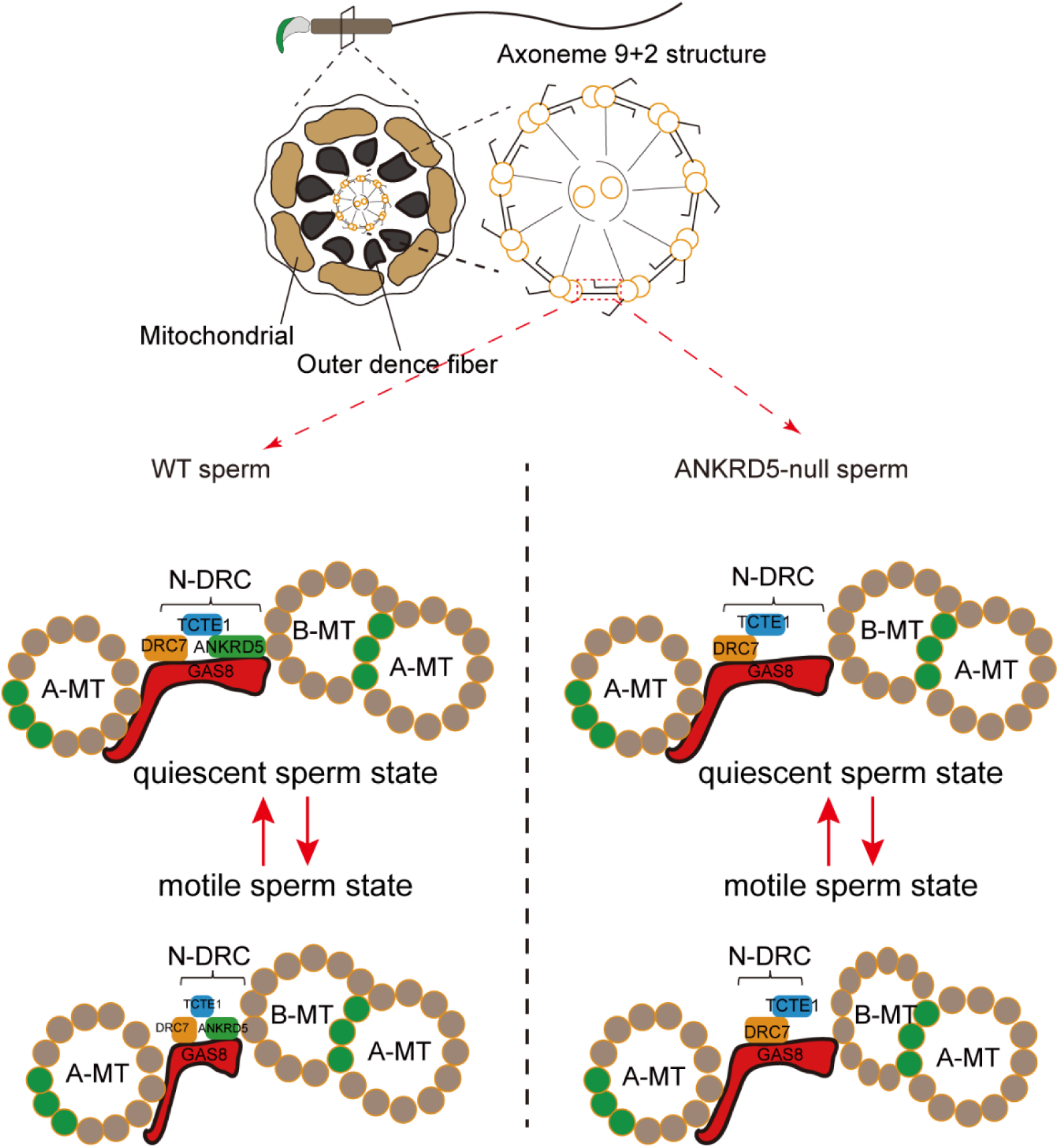

**Significance Statement:** Male infertility affects approximately 8%-12% of men globally, with defects in sperm motility accounting for over 80% of these cases. The axoneme, which functions as the motor apparatus of the sperm, adopts a canonical "9+2" microtubule arrangement, where the nexin-dynein regulatory complex (N-DRC) plays a critical role in providing structural support between adjacent outer microtubule doublets. Elucidating the interplay between the structural organization and protein composition of the N-DRC is essential for advancing the understanding of male reproductive biology. In this study, we identify ANKRD5 as a new component of N-DRC that is essential for maintaining normal sperm motility. These findings contribute to the molecular understanding of sperm motility and highlight ANKRD5 as a potential target for the development of novel male contraceptives.

## Introduction

The interaction between sperm and egg, culminating in embryo formation, is fundamental to sexual reproduction and the continuation of species[1]. Male infertility affects approximately 8% to 12% of the global male population, with defects in sperm motility accounting for over 80% of these cases[2, 3]. Fertilization requires successful spermatogenesis and normal sperm motility[4]. In mammals, sperm acquire motility and fertilizing capacity during transit through the epididymis[5]. This maturation process is essential for generating functionally competent sperm.

Asthenozoospermia, characterized by reduced sperm motility, is a leading cause of clinical infertility; however, its underlying mechanisms remain poorly understood[6]. Men with poorly motile or immobile sperm are typically infertile unless assisted reproductive techniques (ART), such as gamete intrafallopian transfer (GIFT), in vitro fertilization (IVF), or intracytoplasmic sperm injection (ICSI), are employed[7]. Nevertheless, these ART methods may transmit underlying genetic defects to offspring. Deeper insights into the molecular mechanisms of sperm motility could yield targeted therapies for asthenozoospermia. Rather than bypassing the defect with ICSI, such strategies could directly correct it via modulation of key signaling pathways or gene therapy, potentially offering a cure[8, 9, 10].

Sperm motility is powered by the rhythmic beating of the flagellar, which is subdivided into the midpiece, principal piece, and endpiece[11]. These segments share a conserved core structure—the central axoneme—comprising ∼250 proteins that form the main components of the flagellum [12]. The axoneme exhibits a characteristic “9+2” ultrastructure, featuring nine outer doublet microtubules (DMTs) encircling a central pair of singlet microtubules. Adjacent DMTs are interconnected by the N-DRC[13]. The structure and molecular composition of the N-DRC are evolutionarily conserved and central to the regulation of sperm motility[14, 15, 16].

The N-DRC is a ∼1.5 MDa macromolecular complex composed of two primary subdomains: the linker and the base plate[15, 16, 17]. It also interacts with the outer dynein arms (ODA) via outer-inner dynein (OID) linkers, thereby contributing to the regulation of both ODAs and inner dynein arms (IDAs)[18]. Although the N-DRC was initially believed to consist of 11 protein subunits[16, 19], a twelfth component, CCDC153 (DRC12), was later identified through its interaction with DRC1[20]. In situ cryoelectron tomography (cryo-ET) studies in *Chlamydomonas* have elucidated the three-dimensional architecture of the N-DRC, revealing that DRC1, DRC2/CCDC65, and DRC4/GAS8 form the core scaffold[21]. Proteins DRC3/5/6/7/8/11 associate with this core and mediate interactions with other axonemal complexes[22]. Biochemical analyses corroborate these findings and validate the proposed structural model[15, 23, 24]. Functionally positioned between DMTs, the N-DRC converts microtubule sliding into coordinated axonemal bending by restricting the relative displacement of outer DMTs[25, 26, 27]. Genetic mutations in N-DRC subunits demonstrate that its structural integrity is crucial for sperm motility. Specifically, mutations in DRC1, DRC2/CCDC65, and DRC4/GAS8 are associated with ciliary motility disorders, leading to primary ciliary dyskinesia (PCD)[15, 24]. Biallelic truncating mutations in DRC1 induce MMAF in humans, including disassembly of outer DMTs, disorganization of the mitochondrial sheath, and incomplete axonemal assembly[24, 28, 29]. Similarly, loss of CCDC65 destabilizes the N-DRC, resulting in disorganized axonemes, global microtubule dissociation, and complete asthenozoospermia[15, 30]. Recent mammalian knockout studies further confirmed that loss of DRC2 or DRC4 results in severe sperm flagellar assembly defects, multiple morphological abnormalities of the sperm flagella (MMAF), and complete male infertility, highlighting their indispensable roles in spermatogenesis and reproduction[31]. Homozygous frameshift mutations in DRC3 impair N-DRC assembly and intraflagellar transport (IFT), causing severe motility defects despite normal sperm morphology[32, 33]. In contrast, TCTE1 knockout mice exhibit normal sperm axoneme structure but impaired glycolysis, leading to reduced ATP levels, diminished sperm motility, and male infertility[34]. Both *Drc7* and *Iqcg (Drc9)* knockout mice display disrupted "9+2" axonemal architecture, complete sperm immotility, and male infertility[23, 35]. Although the N-DRC is critical for sperm motility, whether additional regulatory components coordinate its function remains unclear. Here, we demonstrate that ANKRD5 is a novel N-DRC component essential for maintaining sperm motility. Absence of ANKRD5 results in diminished sperm motility and consequent male infertility.

## Results

### *Ankrd5* is essential for male fertility

Based on NCBI and single-cell RNA sequencing data, *Ankrd5* exhibits testis-specific expression, with particularly high enrichment in the male reproductive system[36]. In mice, ANKRD5 is a protein of 775 amino acids with a molecular weight of 86.9 kDa. Cross-species sequence comparison revealed that ANKRD5 is evolutionarily conserved (Fig. S1A), and alignment via Clustal Omega demonstrated 86% similarity between mouse and human sequences (Fig. S1B). Quantitative PCR confirmed testis-specific expression of *Ankrd5*, with no detectable expression in brain, liver, spleen, kidney, ovary, intestine, or stomach (Fig. 1A). Temporal expression profiling further revealed that *Ankrd5* expression begins at postnatal day 21 and reaches a relatively high level around day 35 (Fig. 1B), coinciding with the onset of sperm maturation.

**Figure 1.**
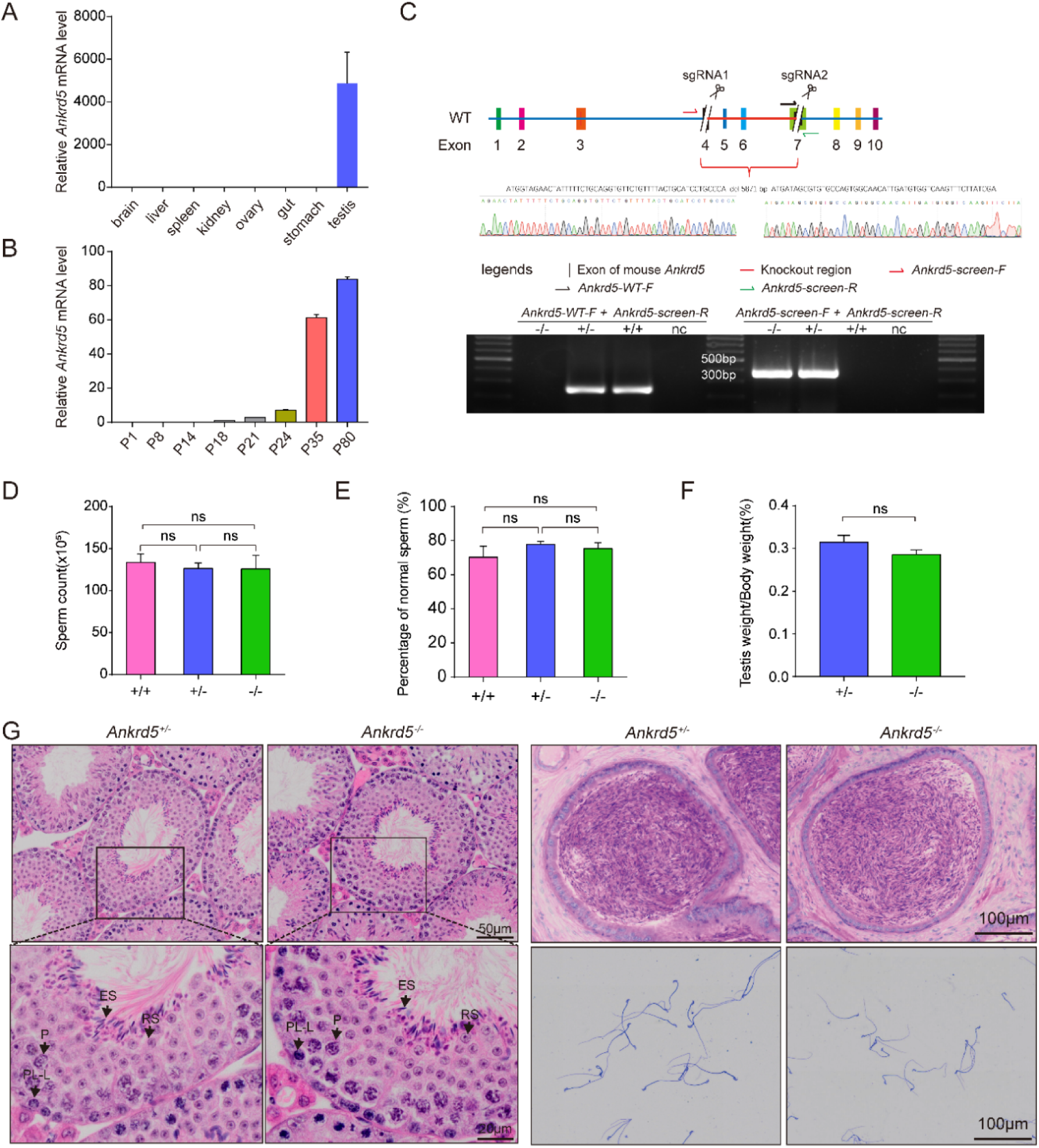
*Ankrd5* is critical for male reproductive function. (A and B) Relative expression of *Ankrd5* mRNA in different tissues of adult mouse and testes at various postnatal days. The *Ankrd5* mRNA expression levels were normalized to the expression of *Gapdh* mRNA (n=3). (C) CRISPR/Cas9 targeting scheme of mouse *Ankrd5* and genotyping of *Ankrd5* KO mouse. *Ankrd5- WT-F* + *Ankrd5-screen-R* (for WT) and *Ankrd5-screen-F* + *Ankrd5-screen-R* (for KO). nc, negative control (ddH_2_O). (D and E) Sperm count and percentage of normal sperm of cauda epididymal from control and *Ankrd5* KO mouse (n=5). (F) Testis to body weight ratio of adult control and *Ankrd5* KO mouse (n=7). (G) Hematoxylin and eosin (H&E) staining of mouse testis and epididymis. Coomassie Brilliant Blue R-250 staining of spermatozoa from control and *Ankrd5* KO male mouse. No significant abnormality was found in Ankrd5 KO male mouse. No overt abnormalities were found in *Ankrd5* KO mouse. P, pachytene; ES, elongated sperm; RS, round sperm; SG, spermatogonia; ST, Sertoli cell. All values in this figure are shown as the meanD±DSE.

To investigate its function in vivo, we generated *Ankrd5* knockout mice (*Ankrd5^−/−^*) on a C57BL/6J background by deleting exons 4-7 using CRISPR/Cas9. Gene deletion was confirmed by PCR(Fig. 1C). Fertility testing revealed that *Ankrd5^−/−^* males were capable of mating with wild-type females but failed to produce offspring (Table 1), indicating complete male infertility. Since litter size and spermatogenesis were unaffected in *Ankrd5^+/–^* males (Table 1; Fig. 1 D-E), heterozygotes were used as experimental controls.

**Table 1.**
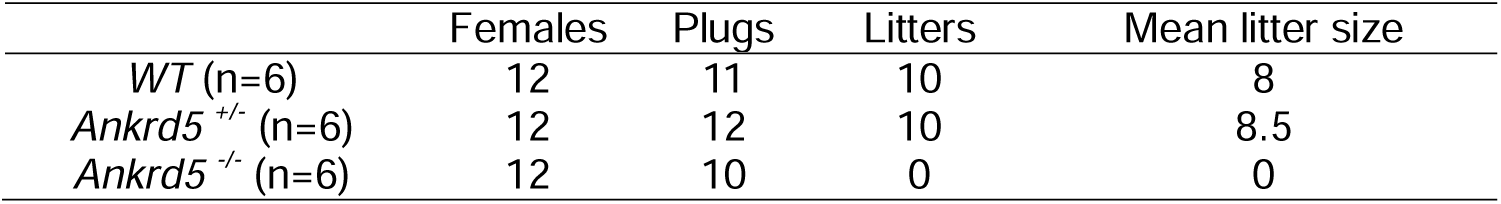
Knockout of *Ankrd5* causes male infertility of mouse Male mouse.

|  | Females | Plugs | Litters | Mean litter size |
| --- | --- | --- | --- | --- |
| *WT* (n=6) | 12 | 11 | 10 | 8 |
| *Ankrd5*<sup>+/-</sup> (n=6) | 12 | 12 | 10 | 8.5 |
| *Ankrd5*<sup>-/-</sup> (n=6) | 12 | 10 | 0 | 0 |

Histological analysis revealed no detectable differences in spermatogenesis between *Ankrd5^−/−^* and control males (Fig. 1 DE and G). The testis-to-body weight ratio and overall morphology of the reproductive system were also normal in *Ankrd5^−/−^* mice (Fig. 1F and S1C-D). Furthermore, H&E staining of testis sections showed that the seminiferous tubules in *Ankrd5^−/−^* mice maintained normal architecture and germ cell composition (Fig. 1G). These findings suggest that male infertility in *Ankrd5^−/−^*mice is not a result of defective spermatogenesis, but instead may reflect a functional defect downstream of sperm development.

### Reduced sperm motility in *Ankrd5* knockout mice impairs zona pellucida penetration

To determine the cause of infertility in *Ankrd5* knockout mice, we performed in vitro fertilization (IVF) assays. Control sperm successfully fertilized both cumulus-intact and cumulus-free oocytes (Fig. 2A, B). In contrast, *Ankrd5* null sperm failed to fertilize cumulus-intact oocytes, despite exhibiting normal binding to the zona pellucida (ZP) (Fig. 2A, D). However, when the ZP was removed, *Ankrd5* null sperm were able to fertilize the oocytes and support development to the blastocyst stage (Fig. 2C). These findings suggest that infertility in *Ankrd5* knockout males is primarily due to impaired sperm penetration of the zona pellucida.

**Figure 2.**
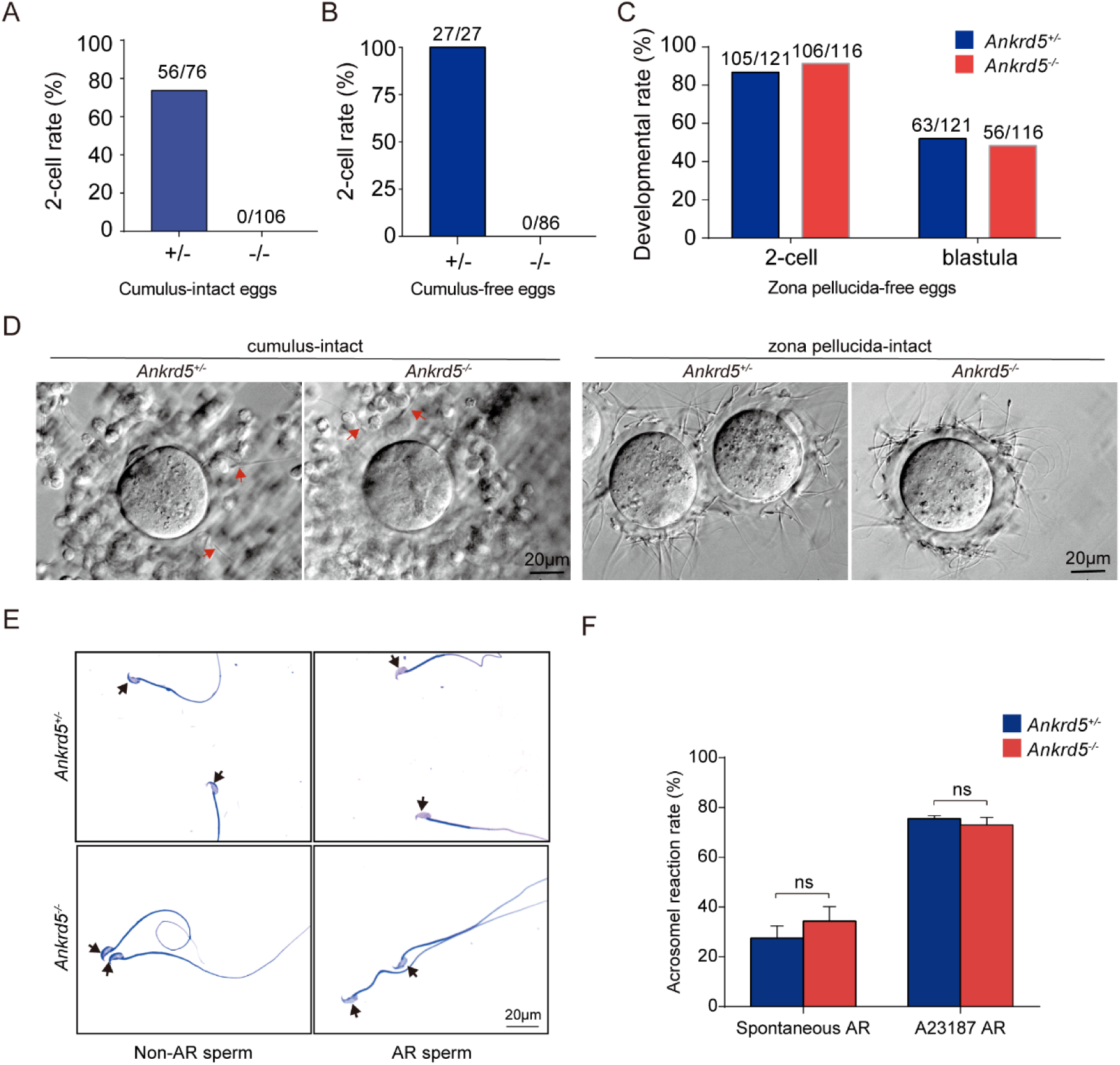
Evaluation of in vitro fertilization capacity of *Ankrd5* KO sperm. (A-C) Fertilization rate of IVF using control and *Ankrd5* KO spermatozoa. Three types of oocytes (cumulus-intact, cumulus-free, and zona pellucida-free) were used for IVF. (D) Egg observation after IVF. After 4Dhours of incubation, both of control and *Ankrd5* KO sperm could penetrate cumulus oophorus as indicated by the red arrow and have the ability to bind to the zona pellucida. (E and F) Sperm were incubated in capacitation medium treated with A23187 (dissolved in DMSO) and DMSO (dissolvent control group) and stained with Coomassie Brilliant Blue R-250. Black arrow indicates the intact or disappeared acrosome; Values represent mean ± SE (n=3).

Penetration through the cumulus-oocyte complex (COC) requires both functional acrosome reaction and adequate motility[37]. Notably, *Ankrd5* null sperm exhibited a normal acrosome reaction when stimulated with the calcium ionophore A23187 (Fig. 2 E, F), indicating that impaired fertilization is not due to a defect in the acrosome reaction.

Since sperm motility plays a vital role in enabling sperm to traverse the cumulus cell layer and the ZP, we next assessed motility using computer-assisted sperm analysis (CASA). Compared with control sperm, *Ankrd5* null sperm showed significantly reduced curvilinear velocity (VCL), average path velocity (VAP), and straight-line velocity (VSL) (Fig. 3A). In addition, parameters reflecting forward progression, including straightness (STR) and linearity (LIN) were also markedly decreased (Fig. 3A). Based on motility classifications (rapid, medium, slow, and static), *Ankrd5* null sperm exhibited a significant decrease in the proportion of rapid sperm and an increase in slow and static subgroups, while the medium subgroup remained unchanged (Fig. 3B). Both total and progressive motility were significantly impaired in the absence of ANKRD5 (Fig. 3C). Time-lapse tracking revealed that *Ankr5* null sperm displayed disorganized and limited motility compared to control sperm (Fig. 3D).

**Figure 3.**
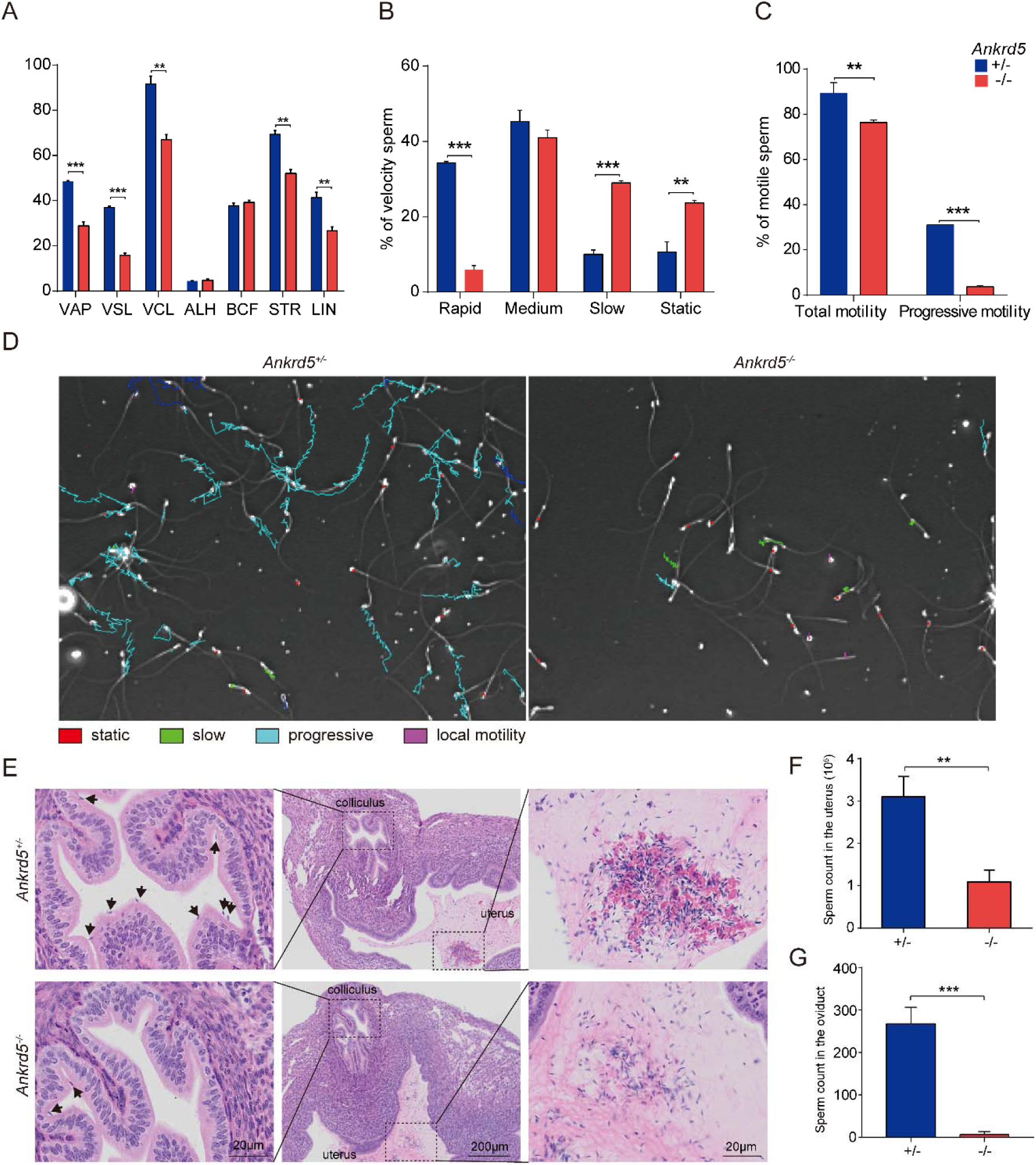
Sperm motility of *Ankrd5* KO male mouse. (A) Average path velocity (VAP), straight line velocity (VSL), curvilinear velocity (VCL), amplitude of lateral head displacement (ALH), beat cross frequency (BCF), straightness (STR), and linearity (LIN) of sperm from control and *Ankrd5* KO mouse. \*\**P* < 0.01, \*\*\**P* < 0.001, Student’s t test; error bars represent SE (n = 3). (B) Proportions of sperm at different velocity levels in control and *Ankrd5* KO mouse. \*\**P* < 0.01, \*\*\**P* < 0.001, Student’s t test; error bars represent SE (n = 3). (C) Knockout mouse had lower motile sperm (total motor capacity) and progressive motile sperm (progressive motor capacity) than control. \*\**P* < 0.01, \*\*\**P* < 0.001, Student’s t test; error bars represent SE (n = 3). (D) Trajectories of sperm per second. The meanings of different colors are shown in the graph. (E) Impaired migration of *Ankrd5* KO sperm from uterus into oviducts. The black arrow indicates sperm. (F and G) Numbers of sperm from control and *Ankrd5* KO mouse in female uterus and oviducts after mating. \*\**P* < 0.01, \*\*\**P* <0.001, Student’s t test; error bars represent SE (n = 3).

To further assess sperm movement in vivo, we tracked sperm trajectories and analyzed their distribution within the female reproductive tract. Sperm migration analysis performed six hours after mating showed a markedly reduced number of *Ankrd5* null sperm in the uterus and oviduct (Fig. 3E-G). These results provide compelling evidence that impaired motility hinders *Ankrd5* null sperm from effectively migrating toward and penetrating the egg, thereby contributing to male infertility.

### ANKRD5 localizes to the sperm axoneme and tracheal motile cilia

To investigate the biological function of ANKRD5, we generated *Ankrd5-*Flag knock-in mice by inserting a Flag tag at the C-terminus via homologous recombination (Fig. S2A). Histological analysis of tissue sections stained with hematoxylin and eosin demonstrated that the epididymal structure and cellular composition in *Ankrd5-*Flag male mice were comparable to those of wild-type controls (Fig. S2B, C). Furthermore, *Ankrd5*-Flag males displayed normal reproductive capacity, indicating that Flag tagging did not disrupt the physiological function of ANKRD5.

Western blot analysis was performed to assess ANKRD5 protein expression across multiple tissues, including liver, lung, kidney, heart, testis, spleen, brain, small intestine, skin, and sperm. ANKRD5 was found to be highly expressed in sperm, with markedly lower levels observed in the testis (Fig. 4A). To further determine its subcellular localization within sperm, we separated sperm heads and tails via repeated freeze-thaw cycles followed by density gradient centrifugation. Western blot analysis revealed that ANKRD5 was predominantly localized to the sperm tail (Fig. 4B).

**Figure 4.**
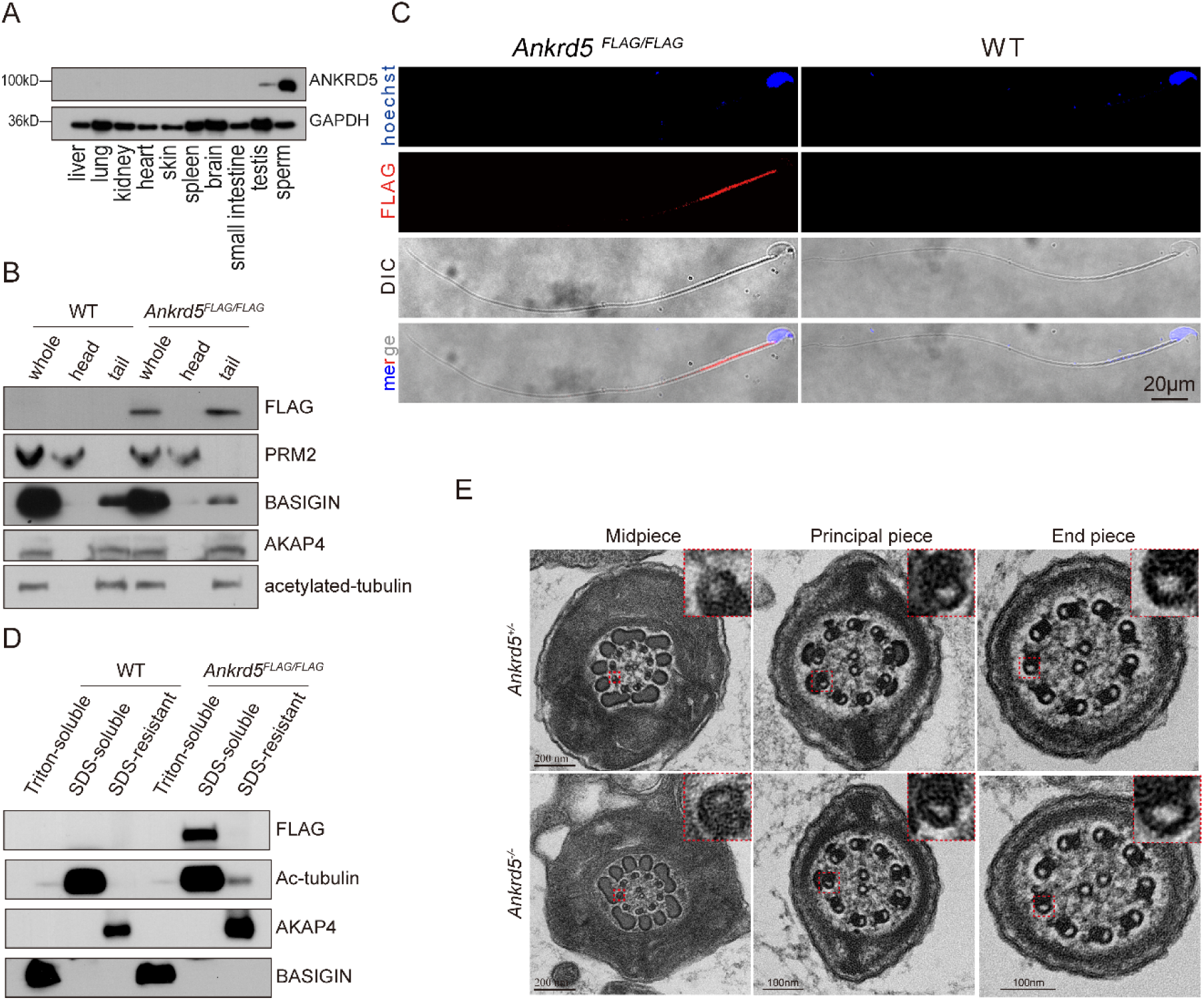
ANKRD5 is located in the midpiece of sperm axoneme. (A) Immunoblotting analysis of various mouse tissues. GAPDH was used as the loading control. (B) Head and tail separation of mouse spermatozoa. ANKRD5-Flag was detected in the tail fraction. PRM2 was used as a marker for sperm head. BASIGIN, AKAP4 and acetylated tubulin were detected as a marker for tails. (C) Immunofluorescence staining results of spermatozoa from wild-type and ANKRD5-Flag mouse using anti-Flag antibody (red: anti-Flag signal; Hoechst: blue). (D) Fractionation of sperm proteins using different lysis buffers. ANKRD5-Flag was found in the SDS-soluble fraction that contains axonemal proteins. BASIGIN, acetylated tubulin, and AKAP4 were detected as a marker for Triton-soluble, SDS-soluble, and SDS-resistant fractions, respectively. (E) Transmission electron microscopy (TEM) of sperm tails from control and *Ankrd5* KO mice. Cross-sections of the midpiece, principal piece, and end piece were examined. Red dashed boxes highlight regions of interest, and the magnified views of these boxed areas are shown in the upper right corner of each image. In three independent experiments, 20 sperm cross-sections per mouse were analyzed for each group, with consistent results observed.

The sperm tail is anatomically divided into three segments: the midpiece, principal piece, and endpiece. Immunofluorescence staining of mature sperm showed that ANKRD5 is primarily localized to the midpiece region (Fig. 4C). As previously reported, proteins associated with different substructures of the sperm tail can be extracted using detergents of varying solubilizing strengths[34, 38, 39]. To refine the localization of ANKRD5, we employed biochemical fractionation using established markers: BASIGIN (Triton X-100–soluble), Acetylated Tubulin (SDS–soluble), and AKAP4 (SDS–resistant), which respectively represent membrane/cytosolic proteins, axonemal proteins, and fibrous sheath/outer dense fiber components[34, 38, 39]. ANKRD5 was primarily enriched in the SDS-soluble fraction, suggesting its association with axonemal structures (Fig. 4D).

These findings support the hypothesis that the infertility observed in *Ankrd5* knockout males may stem from axoneme-related defects in the sperm tail that impair motility. Given that respiratory cilia in mice also possess a canonical axonemal architecture [40], we further examined ANKRD5 localization in tracheal tissue (Fig. S3). However, *Ankrd5-*deficient mice did not display abnormalities in viability or general locomotor function beyond male infertility, suggesting that ANKRD5 may exert a tissue-specific role, with essential function limited to the male reproductive system.

### ANKRD5 interacts with N-DRC components in the axoneme

To investigate the function of ANKRD5 in the axoneme, we characterized its interactome using liquid chromatography–mass spectrometry (LC-MS) (Fig. 5A). Among the interacting proteins identified, several known components of the N-DRC were detected, including TCTE1, DRC3/LRRC48, DRC7/CCDC135, and DRC4/GAS8. Notably, TCTE1 ranked among the most abundant hits (Fig. 5A). Previous studies have shown that *Tcte1* null sperm maintain a normal DMT structure but exhibit reduced motility[34], a phenotype also observed in *Ankrd5* null sperm. Although ANKRD5 was identified in the TCTE1 LC-MS profile, a direct interaction between these proteins had not been previously validated[34]. It is possible that the interactions of ANKRD5 with DRC3 and DRC7 are indirect rather than direct physical bindings.

**Figure 5.**
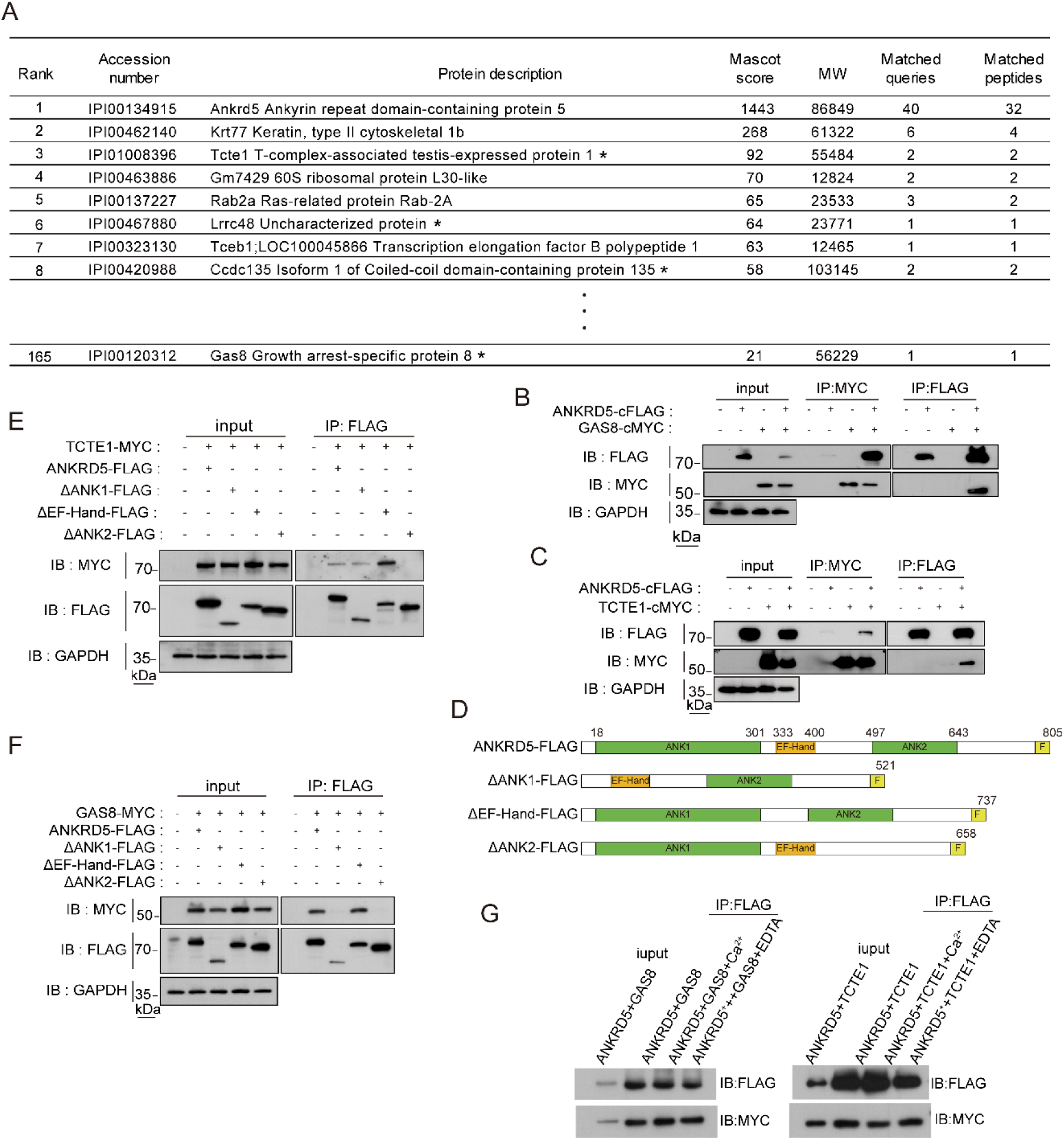
ANKRD5 is a component of N-DRC in sperm flagella. (A) Identification of sperm proteins in LC-MS/MS analysis. Black star indicates N-DRC components. (B and C) Individual DRC components were coexpressed in HEK293T cells. Immunoprecipitation of ANKRD5-Flag resulted in the co-precipitation of GAS8-MYC and TCTE1-MYC. Similarly, immunoprecipitation of GAS8-MYC and TCTE1-MYC also led to the co-precipitation of ANKRD5-Flag. (D) Schematic of various truncated ANKRD5 vectors. Flag-tag was linked posterior to the C-terminal of ANKRD5. Green and yellow boxes show the ANK domain and EF-Hand domain of ANKRD5, respectively. Light yellow boxes indicate Flag tag. (E and F) The interaction between various truncated ANKRD5-Flag and TCTE1-MYC or GAS8-MYC were confirmed by co-IP followed by WB analysis using anti-Flag, and anti-MYC antibodies. (G) Effect of calcium ion and EDTA treatment on the interaction of ANKRD5 with GAS8 and TCTE1.

To confirm these interactions, we performed co-immunoprecipitation assays in HEK293T cells, testing interactions between ANKRD5 and 11 DRC components (excluding DRC6, which failed to express) (Fig. 5B,C; Fig. S4A). These experiments confirmed that ANKRD5 interacts specifically with TCTE1 and DRC4/GAS8, but no with other tested DRC components (Fig. 5B,C; Fig. S4 A,B).

Structurally, ANKRD5 contains two ANK repeat domains and one EF-hand domain. The ANK domain is a well-known scaffold for protein–protein interactions, while the EF-hand domain typically functions as a calcium-binding motif, though approximately one-third EF-hand domains lack calcium-binding ability[41]. Protein truncation analysis revealed that the ANK2 domain is necessary for interaction with TCTE1 (Fig. 5D-E), whereas binding to DRC4/GAS8 requires both ANK1 and ANK2 domains (Fig. 5F). The EF-Hand domain was not essential for interaction with either protein. Furthermore, calcium supplementation did not alter the binding of ANKRD5 to DRC4/GAS8 or TCTE1, as shown by western blot analysis (Fig. 5G), indicating that these interactions are calcium-independent.

Additional candidate interactors identified in the LC-MS dataset include KRT77, a cytoskeletal protein known to maintain structural stability. It may contribute to reinforcing the physical linkage between the N-DRC and adjacent DMTs through interaction with ANKRD5. Recent structural studies have positioned ANKRD5 within the distal lobe of the N-DRC, where its positively charged surface may facilitate electrostatic interactions with glutamylated tubulin on neighboring DMTs[42]. KRT77 may further modulate this interaction via post-translational modifications (PTMs) such as phosphorylation, thereby enhancing the structural integrity of the flagellum under mechanical stress during sperm motility.

Other candidate interactors include members of the Rab GTPase family, which are implicated in intraflagellar transport and membrane trafficking. RAB2A,for example, may regulate the targeted delivery of ANKRD5 or other N-DRC components to assembly sites within the axoneme through vesicle-mediated transport. Its GTPase activity may also serve as a regulatory node linking signaling pathways to axonemal remodeling. However, given the complexity of mass spectrometry datasets, we cannot exclude the possibility that some observed interactions are false positives arising from nonspecific factors such as electrostatic interactions, detergent-mediated membrane disruption, protein aggregation, or high-abundance protein interference.

Sperm motility is highly dependent on energy metabolism. The mitochondrial sheath, located in the sperm tail midpiece, is responsible for ATP production. Previous studies have shown that *Tcte1* null sperm exhibit reduced ATP levels[34]. However, in contrast, *Ankrd5* KO sperm displayed no significant change in ATP content (Fig. 6E), suggesting that ANKRD5 functions independently of ATP synthesis.

**Figure 6.**
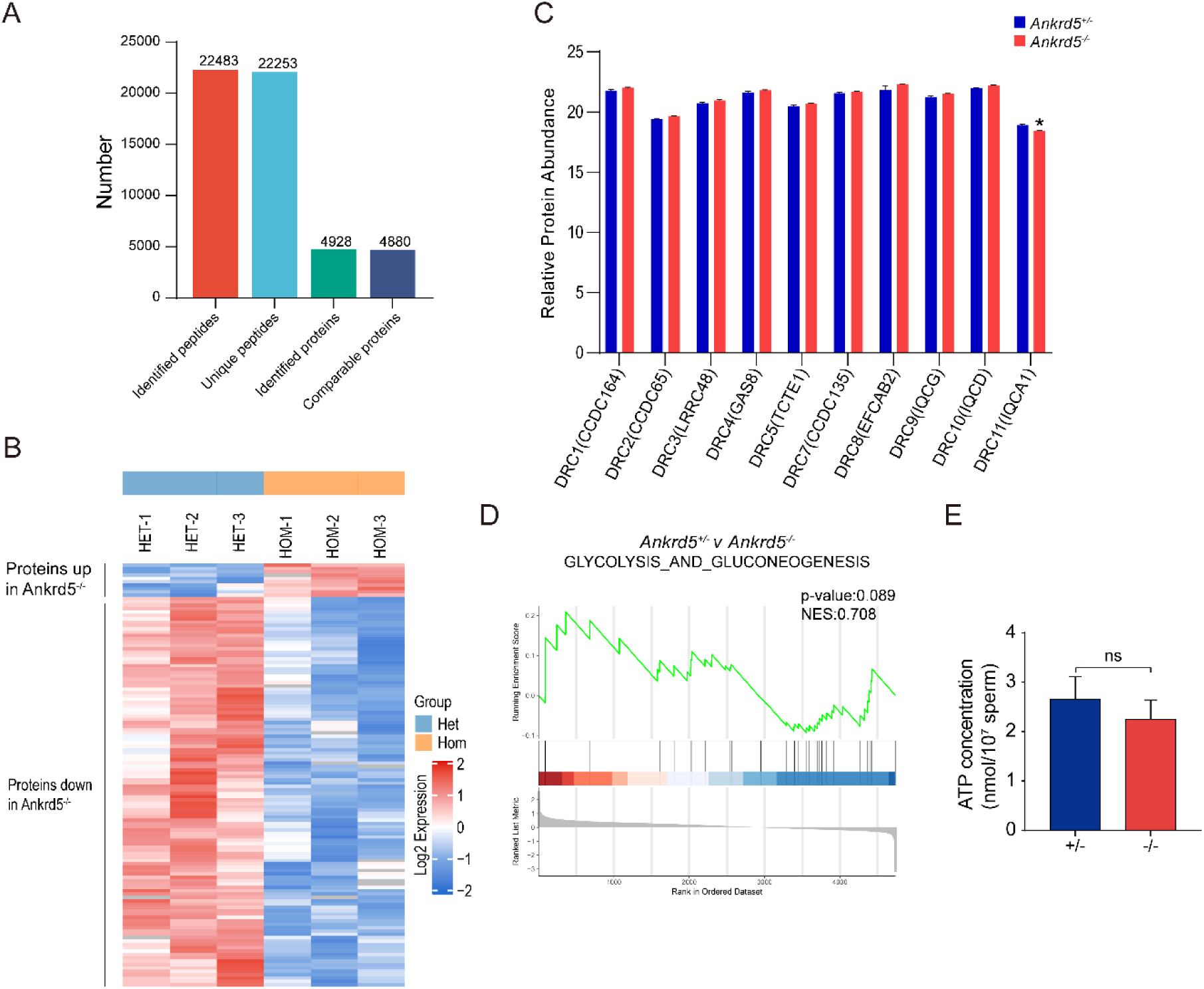
Absence of ANKRD5 does not affect energy metabolism. (A) The differentially expressed proteins of *Ankrd5^+/–^* and *Ankrd5^-/-^* were identified by 4D-SmartDIA. (B) Heatmap of relative protein abundance changes between control and knockout mouse sperm. (C) Differences in the expression of N-DRC protein components identified by mass spectra. \**P* < 0.05, Student’s t test; error bars represent SE (n = 3). (D) GSEA analysis of glycolysis and gluconeogenesis. (E) Measured levels of ATP between wild-type and *Ankrd5* null sperm. Student’s t test; error bars represent SE (n = 3).

The sperm midpiece is also a major site of reactive oxygen species (ROS) generation[43, 44]. Proper regulation of ROS is essential for key sperm functions, including motility, capacitation, acrosome reaction, fertilization, and hyperactivation[45, 46, 47]. Mitochondrial membrane potential (MMP) is correlated with sperm motility[48], decreased MMP (depolarization) indicates mitochondrial dysfunction, while increased MMP (hyperpolarization) leads to excess ROS. We assessed MMP in spermatozoa using TMRM staining and fluorescence imaging (Fig. S5A, B) and measured ROS levels using DCFH-DA staining followed by fluorescence imaging (Fig. S5C, D). No significant differences were observed between *Ankrd5* null sperm and controls, indicating that ANKRD5 deficiency does not significantly affect sperm ATP levels or mitochondrial function.

### The ANKRD5-KO sperm axoneme retains the canonical **"**9+2**"** architecture

Loss of certain DRC components has been shown to affect the expression of other N-DRC subunits. For example, expression of DRC2/3/4 is reduced in *Drc1* mutant mice[24], while DRC1/3/5/11 expression is downregulated in *Chlamydomonas Drc2* mutants[15]. To evaluate the impact of ANKRD5 deletion on sperm protein composition, we performed quantitative proteomic profiling using 4D-SmartDIA. Among the 4,880 quantifiable proteins detected by mass spectrometry, 126 were differentially expressed by more than 1.5 fold, with 10 upregulated and 116 downregulated proteins (Fig. 6A, B).

Notably, none of the annotated N-DRC components were included in the list of significantly differentially expressed proteins. To further examine expression changes of specific N-DRC components, their intensity values were extracted, normalized, and subjected to t-test analysis. Due to technical limitations, DRC6 and DRC12 were not detected. However, DRC11 (IQCA1) displayed a statistically significant change in expression, although validation by Western blot was not possible due to the lack of an commercial antibody (Fig.6C).

Given the reported link between TCTE1 deficiency, glycolytic defects, and reduced sperm motility through lowered ATP levels[34], we conducted Gene Set Enrichment Analysis (GSEA) focusing on glycolysis-related pathways. The analysis yielded no significant enrichment, which aligns with our direct ATP quantification results, indicating that *Ankrd5* knockout does not impair ATP production in sperm (Fig.6D, E). To further evaluate structural integrity of key axonemal elements, including RS, IDA, and ODA, immunofluorescence staining was performed on spermatozoa from *Ankrd5^-/-^* and control mice. No discernible differences were observed (Fig. S6).

The sperm axoneme is organized in a characteristic "9+2" arrangement, consisting of nine outer DMTs surrounding a central pair of singlet microtubules. Structural defects in this 9+2 arrangement may impair motility, but transmission electron microscopy of sperm flagella did not reveal any structural defects in the overall "9+2" structure of the *Ankrd5^-/-^* mice axonemes (Fig. 4E).

To further investigate the potential effects of *Ankrd5* deficiency on axonemal architecture, we collected approximately 160 tilt series of *Ankrd5* null sperm axoneme using cryo-focused ion beam (cryo-FIB) milling followed by cryo-electron tomography (cryo-ET) (Fig. S7). Data pre-processing and reconstruction were performed by Warp and AreTOMO[49, 50, 51] (Fig. S8). In the original tomograms obtained, we found that the overall "9+2" structure of sperm axonemes remained intact regardless of side view or top view, and there was no significant difference from that of WT mouse sperm axonemes (Fig. 7A)[52], which was similar to TEM results. In addition, the presence of DMT accessory structures RS and ODA was observed in both WT and ANKRD5- KO tomograms (Fig. 7A and Fig. S9).

**Figure 7.**
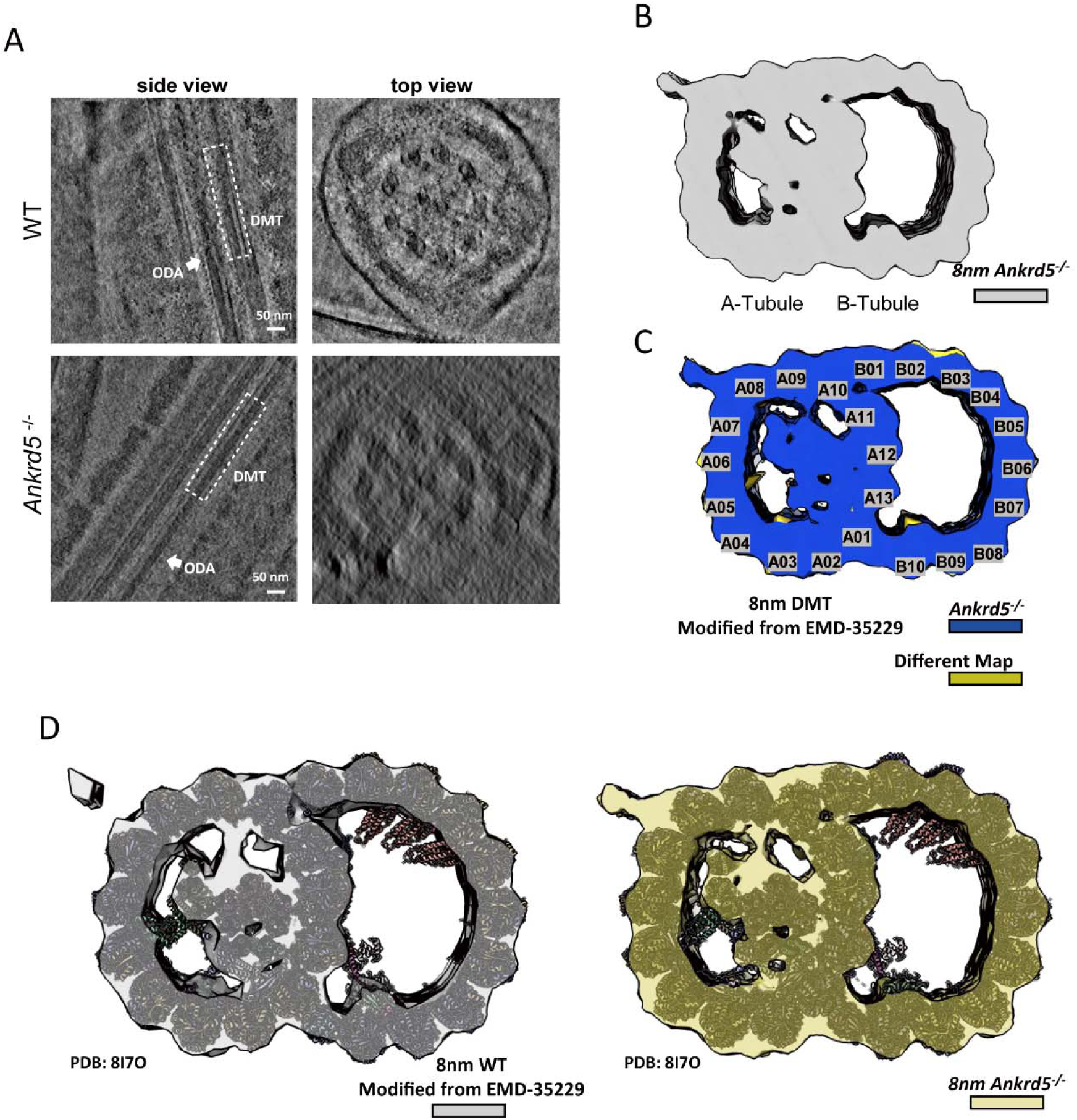
The overall structure of *Ankrd5*-KO mouse sperm DMT. (A) Side view and top sectional view of WT/*Ankrd5*^-/-^ mouse sperm axoneme are shown in the tomogram slices. DMT and ODA are marked with white dashed lines and white arrows, respectively. (B) The cryo-EM map of *Ankrd5*^-/-^ mouse sperm DMT with an 8 nm repeat was obtained by sub-tomogram analysis. (C) Loss of density in *Ankrd5*^-/-^ DMT structure. The transverse sectional view of DMT is shown. The lost density (khaki color) was obtained by subtracting the density map of *Ankrd5*^-/-^ DMT from that of the WT DMT. (D) The model of 16nm-repeats WT DMT (PDB: 8I7O) was fitted in the 8nm repeat WT DMT map and *Ankrd5*^-/-^ DMT map. The 8nm repeats DMT density map was obtained by summing two 16nm repeats DMTs that were staggered 8nm apart from each other.

Among the 89 tomograms with good quality, we carried out manual selection of DMT particles and applied RELION, Dynamo and Warp/M to carry out sub-tomogram averaging (STA) analysis[53, 54, 55]. The 8nm repeat DMT structure of ANKRD5-KO sperm axoneme with a resolution of 24 Å was obtained (Fig. S7B and Fig. S8), and the overall structure of its A and B tubes was complete (Fig. 7 B and D). We compared the DMT density map of ANKRD5-KO mouse sperm with that of WT mouse sperm (Modified from EMD-35229) and found that there was no significant difference between them, except for slight variations in density near A05 of tube A and B10 of tube B (Fig. 7 C and D)[52]. The known mouse sperm DMT model was fitted into two density maps, and all 16nm-repeats MIPs including tubulins could be well fitted. This suggests that the loss of ANKRD5 has no significant effect on the overall structure of axoneme and the component proteins of DMT.

### ANKRD5 depletion may impair the buffering Function between adjacent DMTs in the axoneme

During particle picking of DMT fibers, we observed notable differences in the transverse sections of axonemal DMT particles between ANKRD5-KO and WT sperm. While both A- and B- tubes were discernible in both groups, DMTs in ANKRD5-KO sperm exhibited a markedly more irregular morphology. In WT sperm, DMTs typically appeared circular in cross-section, whereas in ANKRD5-KO sperm, they frequently adopted a polygonal or extruded appearance (Fig. S9B,D). Notably, some ANKRD5-KO DMTs appeared partially open at the A- and B-tubes junctions (Fig. S9B,D).

During the STA process, many ANKRD5-KO particles were either misaligned or displayed significant structural deformation, particularly affecting the B-tube (Figure 9, Fig. S9E). Upon re-examining the TEM data in light of the Cryo-ET findings, similar abnormalities were observed in the TEM images (Fig.4E, Fig. S10B). Notably, both intact and deformed DMT structures were consistently observed in both TEM and STA analyses, with the deformation of the B-tube being more obvious (Fig.4E, Fig. S10). During the STA process, we could retain only ∼10% of the DMT particles to obtain the final density map for ANKRD5-KO sperm (Fig. S9E), compared to ∼70% in the WT dataset as previously reported[52]. The resulting density map from ANKRD5-KO sperm also exhibited peripheral roughness, indicative of substantial structural heterogeneity (Fig. S9E). Even after excluding a large proportion of deformed particles, the final averaged map still presented noticeable artifacts, suggesting that although the overall architecture of the DMTs is preserved, its structural integrity is significantly compromised (Fig. S9E). Furthermore, attempts to resolve the 96-nm repeat structure did not yield clear densities for radial spokes (RSs) (Fig. S9F), suggesting that ANKRD5 deficiency may also impair the stability of accessory structures, such as RSs[25, 26, 27]. In the raw tomograms, RSs in ANKRD5-KO sperm appeared more irregularly arranged compared to those in WT controls (Fig. S9A, C).

Following the submission of this work, ANKRD5 was reported to localize at the head of the N-DRC, interacting simultaneously with DRC11, DRC7, DRC4, and DRC5[42]. These findings are consistent with our in vitro data demonstrating interactions between ANKRD5 and both DRC4 and DRC5 (Fig. 8C-F). However, the aforementioned study utilized isolated and purified DMT preparations, leaving the precise spatial relationship between ANKRD5 and neighboring DMTs unresolved. To address this, We fitted the published structure of ANKRD5(PDB entry: 9FQR) into the in situ 96-nm DMT repeat map from mouse sperm (EMD-27444), revealing that ANKRD5 resides approximately 3 nm from the adjacent DMT (Fig. 8G). The N-DRC is often analogized to a "car bumper", serving as a mechanical buffer between adjacent DMTs during vigorous axonemal motion. Given the pronounced DMT deformation observed in our cryo-ET data (Fig. S9E), we propose that ANKRD5 contributes to this buffering function. Loss of ANKRD5 may compromise the structural resilience of the N-DRC, thereby diminishing its ability to protect adjacent DMTs from mechanical stress and destabilizing associated axonemal accessory structures[20, 42, 56].

**Figure 8.**
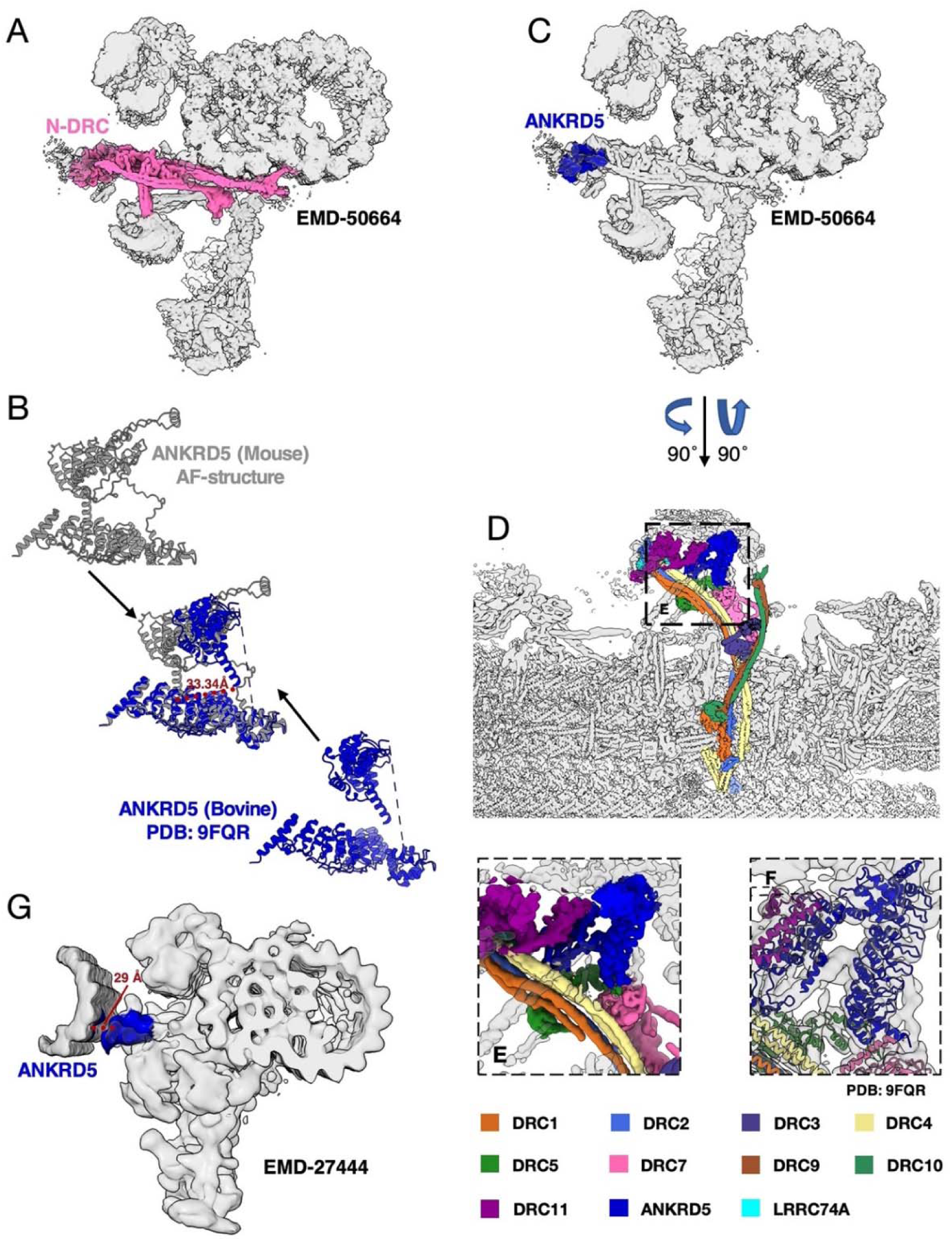
Localization of ANKRD5 in sperm axoneme. (A) Localization of N-DRC in 96nm repeats DMT of mammalian sperm (EMD-50664). (B) Comparison of AlphaFold predicted mouse ANKRD5 structures with known bovine ANKRD5 structures (PDB: 9FQR). (C) Localization of ANKRD5 in 96nm repeats DMT map (EMD-50664). (D) Structure and composition of N-DRC in 96nm repeats DMT of mammalian sperm (EMD-50664). (E) The relationship between ANKRD5 and its interacting proteins as shown in the electron microscope density map (EMD-50664). (F) The relationship between ANKRD5 and its interacting proteins as shown in 96nm repeats DMT model of mammalian sperm (PDB: 9FQR). (G) Localization of ANKRD5 in the in-situ mouse sprem 96nm repeats DMT map (EMD-27444).

**Figure 9.**
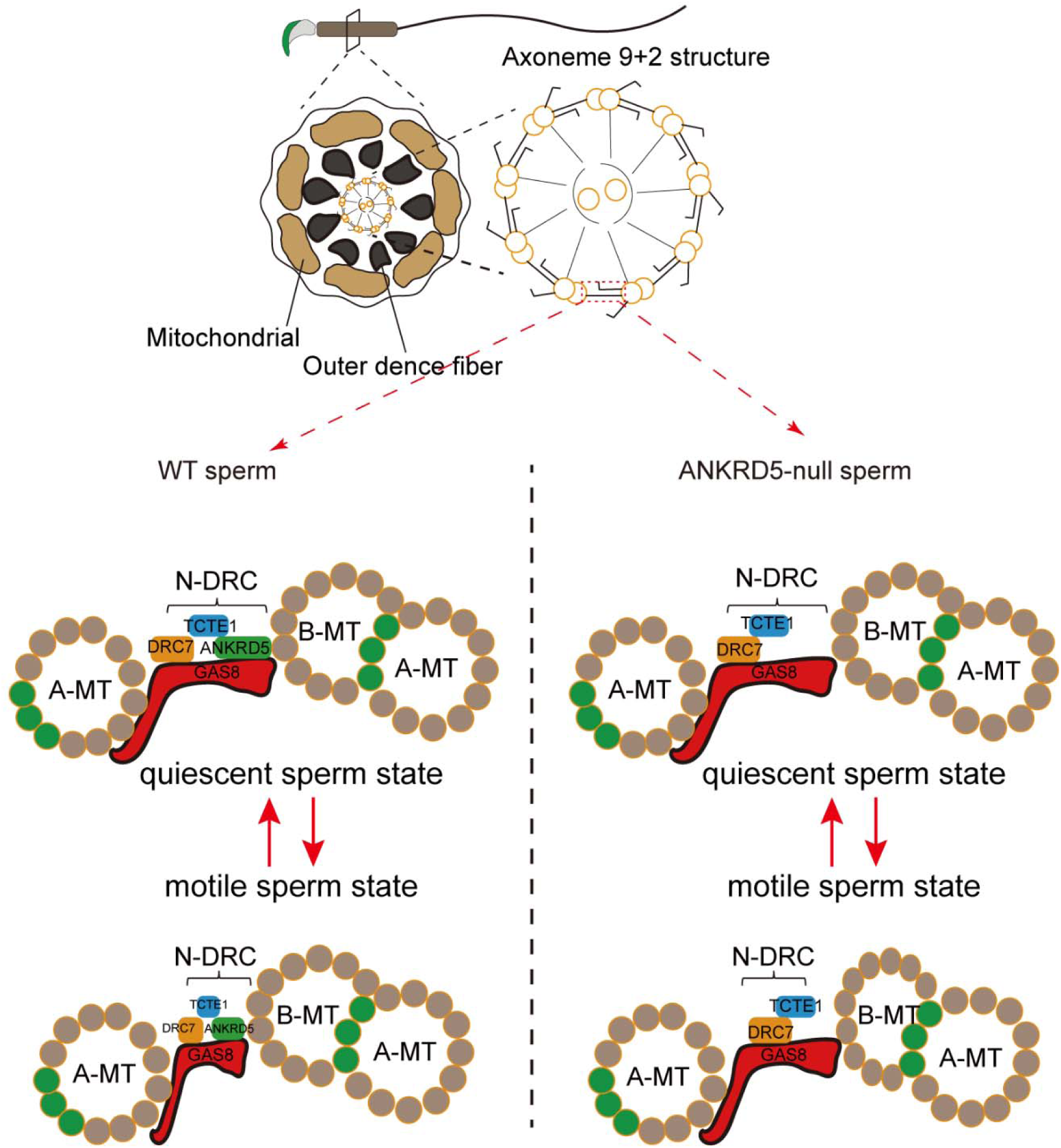
The functional model of ANKRD5. ANKRD5, as a component of the N-DRC connecting adjacent DMTs, enhances the elasticity of the N-DRC. During sperm movement, the flagellar beating causes adjacent DMT to move towards each other, and the intact N-DRC structure provides good cushioning. In ANKRD5 -knockout mice, the absence of ANKRD5 weakens the N-DRC’s buffering capacity. This subjects the axoneme, especially during intense movement, to greater mechanical stress, leading to the deformation of B tube and impaired sperm motility.

## Discussion

During natural fertilization, males ejaculate millions of sperm; however, the majority are expelled by contractions within the female reproductive tract. Freshly ejaculated sperm display activated motility, yet only a few hundred reach the isthmus of the fallopian tube, where they bind to the mucosal epithelium to form a sperm reservoir. In this dormant state, sperm remain quiescent until ovulation occurs [57]. Upon release from the reservoir, sperm undergo hyperactivation, enabling them to approach and penetrate the egg. Activation sperm exhibit symmetrical flagellar beats and swim along near-linear trajectories, whereas hyperactivated sperm show high-amplitude, asymmetrical flagellar movements [58]. Notably, activation is a prerequisite for hyperactivation.

The outer layer of the COC is rich in hyaluronic acid, rendering it highly viscoelastic[59]. During the acrosome reaction, sperm release hyaluronidase, which degrades hyaluronic acid, softening the cumulus matrix and facilitating sperm-egg interaction. Sperm motility is crucial for penetrating the COC[37]. Notably, vole sperm have been reported to penetrate the zona pellucida of mice and hamsters without undergoing an acrosome reaction, highlighting the role of mechanical forces in fertilization [60].

N-DRC is essential for sperm motility. However, whether additional structural components of the N-DRC remain to be identified has not yet been fully determined. As a key architectural element of the axoneme, the proper function of the N-DRC relies on intricate protein-protein interactions. ANKRD5 is an evolutionarily conserved protein characterized by a helix-turn-helix repeat structure known as the ANK domain, a motif widely implicated in mediating protein interactions[61]. Given its high expression in the testis and its identification in the TCTE1- associated LC-MS results, we hypothesized that ANKRD5 may exert a synergistic role in the functional regulation of the N-DRC.

Previous studies have demonstrated that ANKRD5 contributes to cellular adhesion and protrusion, processes critical for the morphogenesis of the Xenopus gastrula[62]. Moreover, ANKRD5 expression differs between normozoospermic and asthenozoospermic males[63], suggesting its potential relevance to sperm function. In our study, *Ankrd5^-/-^*male mice exhibited normal spermatogenesis and successfully mated and produced mating plugs, yet failed to generate offspring. Histological and morphological analyses of testes and epididymal sperm revealed no overt abnormalities. IVF results demonstrated that *Ankrd5* null sperm were capable of penetrating the cumulus cells layer but failed to fertilize cumulus-intact oocytes, even following granulosa cell removal. Strikingly, fertilization was successfully when ZP-free eggs were used, and embryos developed to the blastocyst stage without apparent defects. These findings indicate that ANKRD5 deficiency leads to results in male infertility due to the sperm’s inability to penetrate the ZP.

Penetration of the ZP requires both a functional acrosome reaction and effective sperm motility. Treatment with calcium ionophore A23187 revealed that *Ankrd5* null sperm underwent normal acrosome reactions. However, CASA showed reduced motility, suggesting that impaired ZP penetration is attributable to defective motility rather than a compromised acrosome reaction. This conclusion was further supported by sperm migration assays.

The sperm axoneme, a highly conserved "9+2" microtubule-based structure, is the central driver of sperm motility. Dynein arms anchored to A-tubules generate force through ATP hydrolysis, enabling sliding along adjacent B-tubules. The N-DRC is essential for converting this microtubule sliding into coordinated flagellar bending[25, 26]. Proteomic analysis identified multiple N-DRC components, including DRC5/TCTE1, DRC3/LRRC48, DRC7/CCDC135, and DRC4/GAS8. The sperm phenotype observed in ANKRD5 knockout mice closely mirrors that of TCTE1-deficient mice, suggesting that ANKRD5 may play a functionally analogous role. Supporting this, *Drc3/Lrrc48* knockout mice reach sexual maturity and exhibit normal mating behavior but they are infertile[64]. In chlamydomonas, mutations in *Drc3/Lrrc48* lead to flagellar motility defects[14]. Similarly, *Drc7/Ccdc135* knockout mice display truncated sperm tails and infertility[23], and mutations in DRC4/GAS8 have been implicated in PCD in humans[65].

Our immunoprecipitation experiments confirmed the proteomic interactions identified by mass spectrometry, and GSEA along with ATP quantification demonstrated that ANKRD5 deletion does not impair cellular energy metabolism. These results suggest that ANKRD5 primarily contributes to the structural integrity and function of the N-DRC, independent of metabolic pathways.

Our cryo-ET and TEM results revealed that the loss of ANKRD5 causes deformation of the B-tubule in the sperm axonemal DMT structure, which may be one of the reasons for the reduced sperm motility. Recent elucidation of the 96-nm repeating unit structure of DMT in bovine sperm axonemes has significantly advanced our understanding of the spatial organization of ANKRD5 and its potential interactions with neighboring axonemal proteins[42]. Nevertheless, the structural characterization of N-DRC remains incomplete, particularly regarding its binding interface with adjacent DMTs. High-resolution structural studies at the cellular level are still needed to define the precise architecture of the N-DRC and its connectivity to adjacent axonemal elements. Such insights would contribute to a more comprehensive understanding of the mechanistic basis of mammalian sperm motility.

Asthenospermia, defined by reduced sperm motility, represents a major cause of male infertility, yet its molecular etiology remains poorly understood. Investigating the infertility phenotype observed in *Ankrd5^-/-^* mice may offer new perspectives on the underlying mechanisms of asthenozoospermia. Although current contraceptive options primarily target women, they are frequently associated with undesirable side effects such as dizziness, chest discomfort, decreased libido, and visual disturbances. Consequently, the development of safe and effective male contraceptive has become a pressing objective. Currently, no oral contraceptive is available for men, but our study offers new perspectives for male contraceptive research.

The potential of ANKRD5 as a target for male contraception arises from its essential structural role in the sperm flagellum, mediated via its ANK domain, which facilitates interaction with components of the N-DRC. Structural studies suggest that ANKRD5 features a positively charged surface, which may engage electrostatically with glutamylated tubulin in adjacent microtubules[42], offering a tractable interface for pharmacological targeting. Disruption of this interaction by small-molecule inhibitors could transiently impair sperm motility without affecting systemic physiology. Importantly, sperm function appears to depend more on ANKRD5 than do motile cilia in other tissues, such as those in the respiratory tract, implying that targeted inhibition of ANKRD5 would have minimal impact on ciliary function elsewhere. This type of tissue-specific phenotypic dissociation has been observed in other axonemal proteins, including DNAH17 and IQUB[66, 67], further supporting the feasibility of ANKRD5 as a sperm-specific drug target.

## Materials and Methods

### Animals

All animal experiments were approved by the Animal Care and Use Committee of the Beijing Institute of Biological Sciences. Wild-type C57BL/6 mice were obtained from the SPF Breeding Facility at the Animal Center of the Beijing Institute of Biological Sciences. Animals were housed under a controlled 12-hour light/dark cycle (lights on from 7:00 a.m. to 7:00 p.m.) with ad libitum access to standard chow and water.

### Real-time Quantitative PCR

Total RNA was extracted from specific tissues using TRIzol reagent (Invitrogen, 15596026) according to the manufacturer’s protocol. Tissues were thoroughly homogenized using a mechanical homogenizer and incubated at room temperature for 5 minutes to ensure complete lysis. Chloroform was added and mixed vigorously mixed for 15 seconds, followed by a 3-minute incubation at room temperature. The samples were then centrifuged at 12,000× g for 15 minutes at 4℃ (procedure repeated twice). The aqueous phase was collected, and RNA was precipitated by adding isopropanol, gently inverting the tube several times, and incubating at room temperature for 10 minutes. Samples were centrifuged at 12,000 × g for 10 minutes at 4 °C, and the resulting supernatant was discarded. The RNA pellet was washed with 75% ethanol, vortexed, and centrifuged at 7500 × g for 5 minutes at 4 °C. After removing the ethanol, the pellet was air-dried for 3-5 minutes and resuspended in nuclease-free water. The RNA solution was incubated at 65°C for 3 minutes and then placed immediately on ice. Complementary DNA (cDNA) was synthesized using a reverse transcription kit (TaKaRa, RR047A) following the manufacturer’s instructions. Quantitative PCR (qPCR) was performed using the SYBR Premix Ex Taq kit (TaKaRa, DRR420A), and the relative *Ankrd5* expression levels were normalized to *Gapdh* as the internal control using the ΔΔCt method.

### Generation of *Ankrd5-*deficient and *Ankrd5*-Flag Mouse

Female wild-type mouse aged 4-6 weeks were intraperitoneally injected with pregnant mare serum gonadotropin (PMSG) to stimulate follicular development, followed by injection of human chorionic gonadotropin (hCG, 10 IU/mouse) 47 hours later. After mating with male mice, fertilized zygotes were collected for subsequent microinjection. *Ankrd5^-/-^* mouse were generated by co-injecting Cas9 mRNA with two guide RNAs (sgRNA1 and sgRNA2) into fertilized eggs. The target sequences were as follows: sgRNA1: 5’-cctgcccactaagcggcactatc-3’, sgRNA2: 5’- cctctcatgatagcgtgtgccag-3’. *Ankrd5*-Flag knock-in mouse were generated by co-injecting Cas9 mRNA, a guide RNA, and an *Ankrd5*-Flag plasmid into fertilized embryos. Detailed gene editing strategies are provided in Fig. 1C and Fig.S2A.

### Histological Analysis

Testes and epididymides were dissected and immediately fixed in Davidson’s fixative (formaldehyde: ethanol: glacial acetic acid: distilled water = 6:3:1:10). After an initial fixation at 4 ℃ for 2 hours, the testes were bisected with a razor blade, and tissue fragments were fully immersed in fresh fixative and incubated overnight at 4 °C. The fixed tissues were then dehydrated through a graded ethanol series, embedded in paraffin, and sectioned at a thickness of 5µm. Sections were floated on warm water at 42 °C to flatten, then mounted and baked overnight at 42 °C. Subsequently, the sections were deparaffinized, rehydrated, and stained with hematoxylin and eosin (H&E) following standard protocols. Histological images were acquired using an Olympus VS120 slide scanning microscope.

### In vitro fertilization experiments

Female wild-type mouse aged 4-6 weeks were intraperitoneally injected with pregnant mare serum gonadotropin (PMSG) to stimulate follicular development, followed 47 hours later by human chorionic gonadotropin (hCG, 10IU/mouse) to induce ovulation. Following mating with fertile males, fertilized oocytes were harvested from the females. To remove cumulus cells, oocytes were treated with hyaluronidase (Sigma-Aldrich, H3757) for 10 minutes. For zona pellucida removal, oocytes were treated with Tyrode’s salt solution (Sigma-Aldrich, T1788) for 1 minute. Sperm were introduced into TYH droplets containing either cumulus-free or ZP-free oocytes, at a final concentration of 1x10^6^ sperm/mL. For fertilization assays using cumulus-intact oocytes, the sperm concentration was adjusted to 1x10^4^ sperm/mL. Co-incubation was performed at 37 °C in an atmosphere of 5% CO_2_, the number of embryos reaching the two-cell stage embryos was assessed after 36 hours, and blastocysts formation was evaluated after 3.5 days of culture.

### Acrosome reaction analyses

Sperm were incubated in HTF medium at 37 °C in an atmosphere of 5% CO_2_ for 30 min. A portion of the sperm was treated with the calcium ionophore A23187 (Sigma-Aldrich, 21186-5MG-F) at a final concentration of 10 μmol/L, while the control group was treated with an equivalent volume of DMSO. After 50 minutes of incubation, sperm were fixed with 4% PFA at room temperature for 30 minutes and spread onto glass slides. The slides were then stained with coomassie brilliant blue solution (0.22% coomassie blue R-250, 50% methanol, 10% glacial acetic acid, 40% distilled water) for 10 minutes, followed by rinsing in distilled water to remove excess dye. Images was performed using an Olympus VS120 slide scanning microscope. For each slide, five non-overlapping regions were analyzed, with 100 sperm counted per region.

### Sperm motility analyses

Sperm were incubated in HTF medium at 37 °C in 5% CO_2_ for 1 hour. After removing cauda epididymal tissue, sperm motility was assessed using the CASA system (Version 14 CEROS, Hamilton Thorne Research) equipped with a Slide Warmer (#720230, Hamilton Thorne Research). The acquisition settings included 30 frames captured at a frame rate of 60 Hz. Analysis parameters were as follows: Minimum Contrast: 30, Minimum Cell Size: 4 Pixels and Minimum Static Contrast = 15.

### Sperm Head–Tail Separation

Mouse spermatozoa were subjected to repeated freeze–thaw cycles using liquid nitrogen. The samples were then centrifuged at 10,000× g for 5 minutes, and the resulting pellet was resuspended in 200 μL of PBS. The suspension was gently mixed with 1.8 mL of 90% Percoll solution (sigma, P1644) and centrifuged at 15,000 × g for 15 minutes. Following centrifugation, the sperm heads settled at the bottom of the tube, while tails remained near the top. The separated sperm heads and tails were individually diluted in PBS at a 1:5 volume ratio and centrifuged at 10000 × g for 5 minutes. Each fraction was then washed twice with PBS and lysed using lysis buffer (PH=7.6, 50mM Tris-HCl, 150mM NaCl, 1%TritonX-100, 0.5% sodium deoxycholate, 0.1% SDS, 2mM EDTA) supplemented with protease inhibitor cocktail (Roche, 04693116001). Subsequently, 1/5 volume of 5× loading buffer (PH=6.8,10% SDS, 25% Glycerol, 1M Tris-HCl, 5% β-mercaptoethanol, 0.25% bromophenol blue) was added. The mixture was boiled at 100 °C for 10 minutes, and the supernatant was collected following centrifugation for downstream applications.

### Separation of different sperm fractions

The separation of distinct sperm fractions was performed as previously described[38, 39, 68]. Briefly, sperm collected from the cauda epididymis were washed twice with PBS and resuspended in 1% Triton X-100 lysis buffer (50 mM NaCl, 20 mM Tris ⋅ HCl, pH 7.5) supplemented with a protease inhibitor cocktail. The samples were incubated at 4 °C for 2 hours and subsequently centrifuged at 15,000× g for 10 minutes. The resulting supernatant was designated as the Triton X-100-soluble fraction.

The pellet was then resuspended in 1% SDS lysis buffer (75 mM NaCl, 24 mM EDTA, pH 6.0), incubated at room temperature for 1 hour and centrifuged. The supernatant from this step was collected as the SDS-soluble fraction. The remaining pellet was boiled for 10 minutes in sample buffer (66 mM Tris-HCl, 2% SDS, 10% glycerol, and 0.005% bromophenol blue), followed by centrifugation. The resulting supernatant was designated as the SDS-resistant fraction. Protease inhibitor cocktail was added to both the Triton X-100 and SDS lysis buffers immediately prior to use.

### Transmission electronic microscopy

Transmission electron microscopy (TEM) was conducted at the Centre for Electron Microscopy, National Institute of Biological Sciences, Beijing, according to standard protocols.

Cauda epididymal tissue was excised and fixed overnight at 4℃ in 2.5% glutaraldehyde (Sigma-Aldrich, G5882). Samples were then washed three times with PBS for 20 minutes each, followed by postfixation in 1% Osmium tetroxide for 1 hour at room temperature. After another three PBS washes, the tissues were dehydrated in a graded acetone series using the progressive lowering of temperature method and subsequently embedded in epoxy resin. The embedded blocks were cured in a drying oven at 45℃ for 12 hours and then at 60℃ for 72 hours. Ultrathin sections were prepared and stained with 3% uranyl acetate in 70% methanol/H_2_O, followed by Sato’s lead stain for 2 minutes. Imaging was performed using a TECNAI spirit G2 (FEI) transmission electron microscope at 120 Kv.

### Western blot

Mouse tissue samples were collected and lysed in buffer (PH=7.6; 50mM Tris-HCl, 150mM NaCl, 1%Triton X-100, 0.5% sodium deoxycholate, 0.1% SDS, and 2mM EDTA) supplemented with a protease inhibitor cocktail. Prior to use, the protease inhibitors were added in appropriate proportion. The lysate waere mixed with 1/5 volume of 5× loading buffer (10% SDS, 25% glycerol, 1M Tris-HCl, 5% β-mercaptoethanol, and 0.25% bromophenol blue, pH 6.8), boiled at 100 °C for 10 minutes, and centrifuged. The resulting supernatants were used for SDS-PAGE. Proteins were resolved by SDS-PAGE and transferred onto PVDF membranes. Membranes were blocked with 5% skim milk in TBST (20 mM Tris, 150 mM NaCl, 0.05% Tween-20, pH 7.6) for 1 hour at room temperature. Membranes were incubated with primary antibodies overnight at 4 °C, followed by three washes with TBST (10 minutes each). Afterward, membranes were incubated with HRP-conjugated secondary antibodies at room temperature for 1 hour and washed again three times with TBST. Chemiluminescent signals were developed using ECL reagents (BIO-RAD, 170-5060 or NCM Biotech, P10300B) and detected using XBT X-ray film (Carestream, 6535876).

### Immunofluorescence

Mouse sperm were collected in pre-warmed PBS and incubated at 37 °C for 15 minutes. The cauda epididymis tissue was discarded, and the remaining sperm suspension was centrifuged at 1000x g for 5 minutes. The sperm pellet was fixed in 4% PFA at room temperature for 30 minutes, spread on glass slides, and air-dried. Antigen retrieval was performed using antigen retrieval buffer (10 mM sodium citrate, 0.05% Tween-20, pH 6.0), followed by cooling to room temperature. Samples were then blocked with ADB blocking solution (3% BSA, 0.05% Triton X- 100) for 1 hour at room temperature. Slides were incubated with the primary antibody (anti-Flag M2) overnight at 4 °C and washed three times with PBST (PBS containing 0.1% Tween-20), 5 minutes each wash. Subsequently, slides were incubated for 1 hour at room temperature, protected from light, with secondary antibodies (Alexa Fluor® 546-conjugated donkey anti-mouse IgG and Alexa Fluor® 647-conjugated donkey anti-rabbit IgG) along with Hoechst 33342 (Sigma, B2261). After incubation, samples were washed three times with PBST for 5 minutes each. Images were acquired using a Nikon Structured Illumination Microscope (SIM) confocal system.

### Identification of interacting proteome by LC-MS

Protein bands were carefully excised from the polyacrylamide gel using a clean razor blade, ensuring minimal inclusion of excess background gel. The excised gel bands were cut into into smaller fragments and transferred to 1.5 mL microcentrifuge tubes for further processing. Gel pieces were destained three times using 25 mM ammonium bicarbonate (NHLJHCOLJ) in 50% methanol, with each step lasting 10 minutes. This was followed by three washes with 10% acetic acid in 50% methanol for 1 hour each. The gels were then rinsed twice with distilled water for 20 minutes per rinse to remove residual solvents.

After thorough rinsing, the gel pieces were transferred to 0.5 mL microcentrifuge tubes and dehydrated by adding 100% acetonitrile (ACN). Tubes were gently inverted until the gel pieces turned opaque white. The ACN was removed, and the samples were dried in a SpeedVac under mild heating for 20–30 minutes. For proteolytic digestion, the dried gel fragments were rehydrated with 5–20 μL of trypsin solution (10 ng/μL in 50 mM NH₄HCO₃, pH 8.0) and incubated at 37 °C overnight.

Following digestion, peptides were extracted by incubating the gel pieces in 25–50 μL of 50% ACN with 5% formic acid (FA) for 30–60 minutes under gentle agitation (avoiding vortexing). The resulting supernatant was transferred to a new 0.5 mL tube. A second extraction was performed using 25–50 μL of 75% ACN with 0.1% FA under the same conditions. The supernatants from both extractions steps were combined and dried completely using the SpeedVac under gentle heat to ensure full peptide recovery for downstream LC-MS analysis.

### Verification of protein interactions in 293T cells by co-immunoprecipitation

Candidate protein-coding sequences were obtained from the NCBI database and cloned into the pcDNA3.1 expression vector. The primer sequences used for vector construction are listed in Supplementary Material 1. Plasmids were transfected into HEK293T cell using the jetOPTIMUS® transfection reagent (cat. No: 101000006), and cells were incubated at 37 °C in a 5% COLJ atmosphere for 36 hours.

Following incubation, the culture medium was removed, and the cells were gently washed once with PBS. Cell lysis was performed using RIPA buffer (pH 7.6: 50 mM Tris-HCl, 150 mM NaCl, 1% Triton X-100 or 1% NP-40, 2 mM EDTA), supplemented with protease inhibitors as needed. A portion of the lysate was reserved as the input control, while the remaining sample was equally divided for immunoprecipitation (IP) using either anti-Flag or anti-Myc antibodies. The IP reactions were incubated overnight at 4 °C with gentle rotation.

The immune complexes were captured using protein-conjugated beads, which were washed five times with RIPA buffer (3 minutes per wash). Bound proteins were eluted by adding SDS loading buffer, followed by denaturation at 95 °C for 5 minutes. Eluted samples were subjected to SDS-PAGE and immunoblotting to assess protein–protein interactions.

### Proteomic analysis of whole sperm

Sperm samples were ground into a fine powder using liquid nitrogen and transferred into microcentrifuge tubes. Lysis buffer (8 M urea supplemented with1% protease inhibitor cocktail) was added at a 1:4 ratio. The samples were sonicated on ice for 3 minutes using a Scientz high-intensity ultrasonic processor. Following sonication, the lysates were centrifuged at 12,000 ×g for 10 minutes at 4 °C, and the supernatants were collected. Protein concentrations were determined using a BCA assay kit according to the manufacturer’s protocol.

For protein digestion, the samples were first reduced with 5 mM DTT at 56 °C for 30 minutes, followed by alkylation with 11 mM iodoacetamide in the dark at room temperature for 15 minutes. The solution was then diluted with 200 mM TEAB to reduce the urea concentration to below 2 M. Trypsin was added at 1:50 ratio for overnight digestion, followed by a second digestion at 1:100 ratio for 4 hours to ensure complete proteolysis. Peptides were subsequently purified using Strata X SPE columns.

For LC-MS/MS analysis, the purified peptides were dissolved in solvent A (0.1% formic acid and 2% acetonitrile in water) and separated on a 25 cm in-house packed reversed-phase column (inner diameter: 100 μm). Chromatographic separation was achieved using a linear gradient of 6% to 80% solvent B (0.1% formic acid, 90% acetonitrile) over 20 minutes at a flow rate of 700 nL/min using an EASY-nLC 1200 ultra-performance liquid chromatography (UPLC) system.

Mass spectrometry was performed on an Orbitrap Exploris 480 instrument. Full MS scans were acquired at a resolution of 60,000, and MS/MS spectra were obtained at a resolution of 15,000 using HCD with a 27% NCE. Data-independent acquisition (DIA) data were processed using DIA-NN software. Spectra were searched against the *Mus musculus* UniProt database (Mus_musculus_10090_SP_20231220.fasta), specifying Trypsin/P specified as the digestion enzyme and allowing up to one missed cleavage. Fixed modifications included N-terminal methionine excision and carbamidomethylation of cysteine residues. The false discovery rate (FDR) was controlled at <1% at both the peptide and protein levels.

### Measurement of sperm ATP levels

Mouse sperm were Collected in pre-warmed PBS and incubated at 37 °C for 15 minutes. Cauda epididymal tissue was removed, and the remaining solution was centrifuged at 1000 ×g for 5 minutes. The sperm pellets were washed twice with PBS. After counting, 1×10^7^ sperm cells were seeded into each well of a 96-well plate. Subsequently, 30μL of ATP lysis (Promega, G7570) was added to each well. The plates were shaken for 20 minutes in the dark, and ATP levels were quantified using a MicroPlate Spectrophotometer (TECAN) according to the manufacturer’s instructions.

### Mitochondrial membrane potential assessment

Sperm were collected from the cauda epididymis, and mitochondrial membrane potential was assessed using tetramethylrhodamine methyl ester (TMRM; Invitrogen, I34361)[69]. Briefly, mouse sperm were incubated in pre-warmed PBS at 37 °C for 15 minutes. Following removal of the cauda tissue, the suspension was centrifuged at 300 xg for 5 minutes, and the resulting pellets were washed twice with DPBS. The washed sperm were then incubated with TMRM for 30 minutes at 37℃ in a humidified atmosphere containing 5% CO_2_. After incubation, sperm were washed twice with DPBS to remove excess unbound dye. The pellet was resuspended, mixed with DAPI, and the suspension was spread onto microscope slides. Fluorescence images were acquired using a DAPI filter (excitation: 405 nm) and an RFP filter (excitation: 560nm).

### Evaluation of intracellular ROS levels

Spermatozoa were isolated from the cauda epididymis and incubated pre-warmed TYH medium at 37 °C for 50 minutes, with or without the ROS inducer Rosup. After incubation, samples were centrifuged gently at 300 ×g for 4 minutes and resuspended in 1 mL of DCFDA working solution. The DCFDA mix was prepared by diluting DCFH-DA (Beyotime, S0033S) at a 1:1000 ratio in TYH medium. The sperm suspension was incubated at 37 °C with 5% CO_2_ for 20 minutes, with gentle inversion every 5 minutes to ensure even dye distribution. Following incubation, the samples were centrifuged again at 300 ×g for 4 minutes, and the supernatant was discarded. The sperm pellets were then washed three times with DPBS to eliminate excess extracellular DCFH-DA. The final pellet was resuspended, mixed with DAPI, and mounted onto microscope slides. Fluorescence images were captured using a DAPI filter and an FITC filter.

### Sample preparation of mouse sperm axoneme

Freshly collected sperm were centrifuged at 400 ×g for 5 min at 4 °C using a Thermo Scientific Legend Micro 17 R centrifuge. The resulting pellet from every 100μL of semen was gently resuspended in 100μL of pre-cooled PBS and subsequently diluted 5.5-fold with PBS prior to use. Cryo-EM grids (Quantifoil R3.5/1, Au 200 mesh) were glow-discharged for 60 seconds using a Gatan Solarus system. Sperm samples (3 μL, diluted in PBS) were applied to the grids, incubated for 3–5 seconds under 100% relative humidity at 4 °C to allow absorption, and then the frozen samples were placed in a mixture of ethane and methane cooled to -195 °C. The grids were then stored in liquid nitrogen for subsequent cryo-FIB milling. The cryo-FIB thinning strategy followed protocols described in previous work[52].

### Cryo-ET tilt series acquisition

The grid after cryo-FIB milling, was loaded onto the Autoloader in Titan Krios G3 TEM (Thermofisher Scientific) operating at 300 KV. The microscope was equipped with a Gatan K2 direct electron detector (DED) and a BioQuantum energy filter. Tilt series were acquired at a magnification of ×42,000, yielding a physical pixel size of 3.4 Å in counting mode. Prior to data collection, the sample’s pre-tilt was visually assessed and set to either +10° or -9°, depending on the pre-defined geometry induced during grid loading. The total electron dose per tilt was 3.5 electrons/Å², distributed across 10 frames over a 1.2-second exposure. The tilt range was -66° to +51° for a -9° pre-tilt, and -50° to +67° for a +10° pre-tilt, with 3° increments, resulting in 40 tilts per series and a cumulative dose of approximately 140 electrons/Å². The energy filter slit width was set to 20 eV, and the zero loss peak was recalibrated after the acquisition of each tilt series. The nominal defocus range was set between -1.8 μm and -2.5 μm. All tilt series used in this study were acquired using a beam-image shift-assisted, dose-symmetric acquisition scheme implemented with a custom SerialEM software[70, 71, 72].

### Mouse sperm axoneme *in situ* data processing

After data collection, all fractioned movies were imported into Warp for essential processing, including motion correction, 2-x Fourier segmentation of super-resolution frames, CTF estimation, masking platinum islands or other high-contrast features, and tilt series generation. Subsequently, the tilt series was automatically aligned using AreTOMO[50, 51]. Aligned tilt series were visually inspected in IMOD, and any low-quality frames (such as those blocked by the sample stage or grid bars, containing obvious crystalline ice, or showing obvious jumps) were removed to create new sets of tilt series in Warp. The new tilt series underwent a second round of AreTOMO alignment. Then, low-quality frames were removed again using the same criteria as in the first round. The new tilt series was then subjected to a third round of automatic alignment, continuing the process until no frames remained to be removed. After tilt series alignment, those tilt series with fewer than 30 frames or that failed were not further processed[50, 51]. The alignment parameters of all remaining tilt series were passed back to Warp, and initial tomogram reconstruction was performed in Warp at a pixel size of 27.2 Å, resulting in a total of 160 Bin8 tomograms[50].

Among the total 160 tomograms, we used the filament picking tool in Dynamo to manually pick DMT particles from 89 tomograms. By selecting the starting and ending points of each DMT fiber and separating each cutting point by 8 nm along the fiber axis, we obtained 89 sets of DMT particles[73]. The 3D coordinates and two of the three Euler angles (except for in-plane rotation) were automatically generated by Dynamo and then transferred back to Warp for exporting sub-tomograms[50, 73, 74].

In RELION 3.1, sub-tomograms were refined, using the ABTT package to transform the RELION star file and Dynamo table file and to jointly generate a mask using Dynamo and/or RELION[50, 73, 74]. First, all particles were reconstructed into a box size of 48^3^ voxels with a pixel size of 27.2 Å, and all extracted particles were directly averaged and low-pass filtered at 80 Å to generate an initial reference. Then, 3D classification with K = 1 was performed under the constraints of the first two Euler angles (—sigma_tilt 3 and —sigma_psi 3 in RELION), followed by 3D automatic refinement after 25 iterations. After alignment, the particles were manually cleaned in ChimeraX-1.6[52, 75], and the aligned parameters were transferred back to Warp to export the sub-tomograms with a box size of 84^3^ voxels and a pixel size of 13.6 Å. The particles were automatically refined in RELION. After removing any duplicate particles in Dynamo, the aligned parameters were transferred back to Warp to export the sub-tomograms with a box size of 128^3^ voxels and a pixel size of 6.8 Å. The particles were then automatically refined in RELION, resulting in a final resolution of 24 Å[53, 54, 55].

### GSEA analysis

Protein mass spectrometry was performed using samples from three controls and three ANKRD5 knockout mice. Dropout data were imputed using KNN, followed by log2 normalization and sorting to obtain the final data list. Mouse gene symbols were converted to Entrez IDs using the org.Mm.eg.db annotation package, and GSEA analysis was performed with the clusterProfiler package[76]. The term ’Glycolysis and Gluconeogenesis’ was obtained from KEGG database. Mouse gene sets were retrieved from the MSigDB (Molecular Signature Database) via the msigdbr package, focusing on category C2, which includes various biological processes and pathways. The analysis was conducted in R version 4.4.1 (2024-06-14 ucrt).

### Statistical analysis

All data are presented as the mean ± SEM (n≥3). Statistical analysis was performed using Student’s t-test or one-way ANOVA. GraphPad Prism version 9.4.1 was used for the analysis, and results were considered significant when *P* < 0.05. Adobe Illustrator 2021 was used for image layout.

## Acknowledgments

We thank the Transgenic Animal Center at the National Institute of Biological Sciences, Beijing, for their support in generating and maintaining the transgenic mice. We are also grateful to Professor Fei Sun’s group at the Institute of Biophysics, Chinese Academy of Sciences, for their technical assistance in Cryo-ET. Additionally, we appreciate the staff at the Electron Microscopy Center and the Imaging Center of the National Institute of Biological Sciences, Beijing, for their technical support in imaging. We also extend our thanks to the State Key Laboratory for Animal Biotechnology at the College of Biological Sciences, China Agricultural University, for their help in analyzing mouse sperm motility. This work was supported by the Beijing Natural Science Foundation (JQ24056 to Y.Z.) and the National Natural Science Foundation of China (32471244 to Y.Z.).

## Author Contributions

Fengchao Wang, Yun Zhu, Fei Sun, Shuntai Yu, and Guoliang Yin designed research; Shuntai Yu, Guoliang Yin, Peng Jin, Weilin Zhang, Yingchao Tian, Xiaotong Xu, Tianyu Shao, and Yushan Li performed research; Yun Zhu, Fei Sun, Guoliang Yin contributed Cryo-ET expertise; Shuntai Yu, Guoliang Yin, and Fengchao Wang, Yun Zhu, Fei Sun analyzed data; and Shuntai Yu, Guoliang Yin, and Fengchao Wang wrote the paper.

## Competing Interest Statement

The authors declare no competing interest.

---

**Figure S1.**
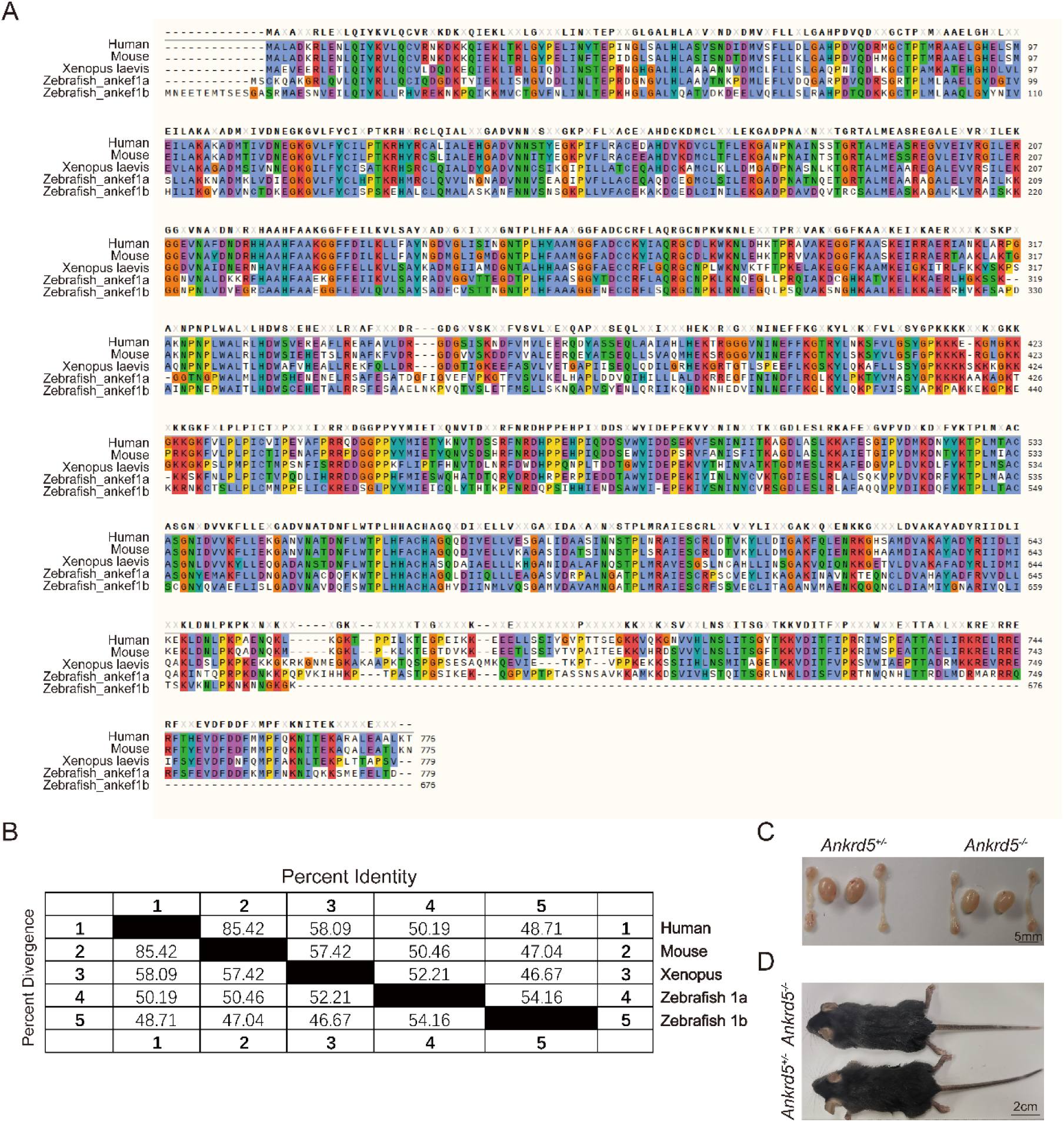
ANKRD5 is conserved between human and other vertebrate model organisms. (A) Sequence alignment of ANKRD5 proteins from several species. Sequences were derived from NCBI and compared with SnapGene (Version=4.3.6). (B) Percent identity matrices of *Ankrd5* proteins across several common vertebrate model organisms. (C) Gross morphology of adult control and *Ankrd5* KO testes and epididymis. (D) The body size was similar between *Ankrd5*^+/-^ and *Ankrd5*^-/-^ mice.

**Figure S2.**
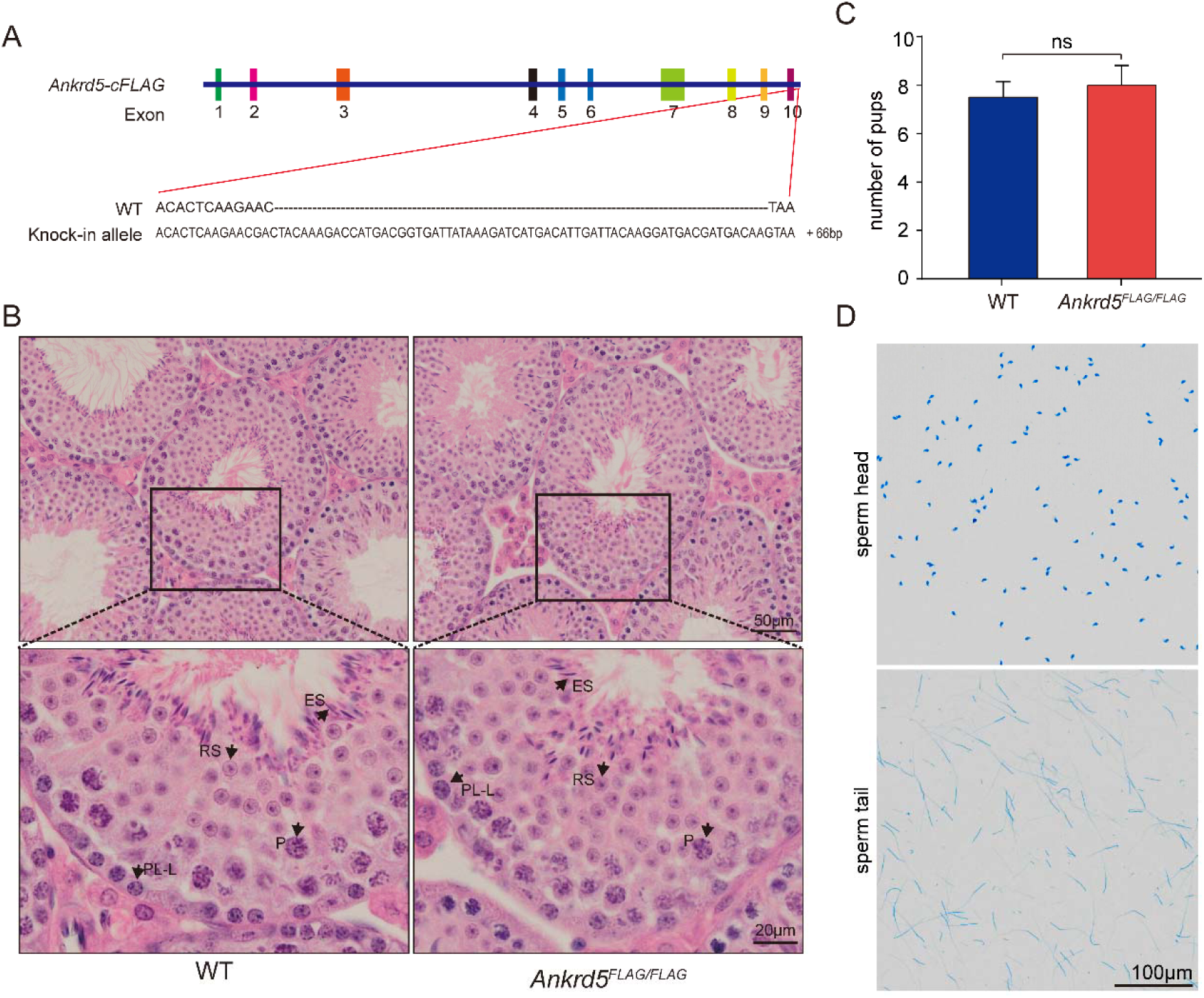
The generation of Flag-tagged mouse and sperm head and tail separation. Schematic of Flag-tagged alleles of endogenous *Ankrd5* generated using CRISPR/Cas9. (B) Hematoxylin and eosin (H&E) staining of Flag-tagged mouse testis. No overt abnormalities were found. P, pachytene; ES, elongated sperm; RS, round sperm; SG, spermatogonia; ST, Sertoli cell. (C) There was no significant difference in litter size between WT and Flag-tagged mouse. Values represent mean ± SE (n=3). (D) Sperm head and tail were separated by repeated freeze-thaw and stained with Coomassie Brilliant Blue R-250.

**Figure S3.**
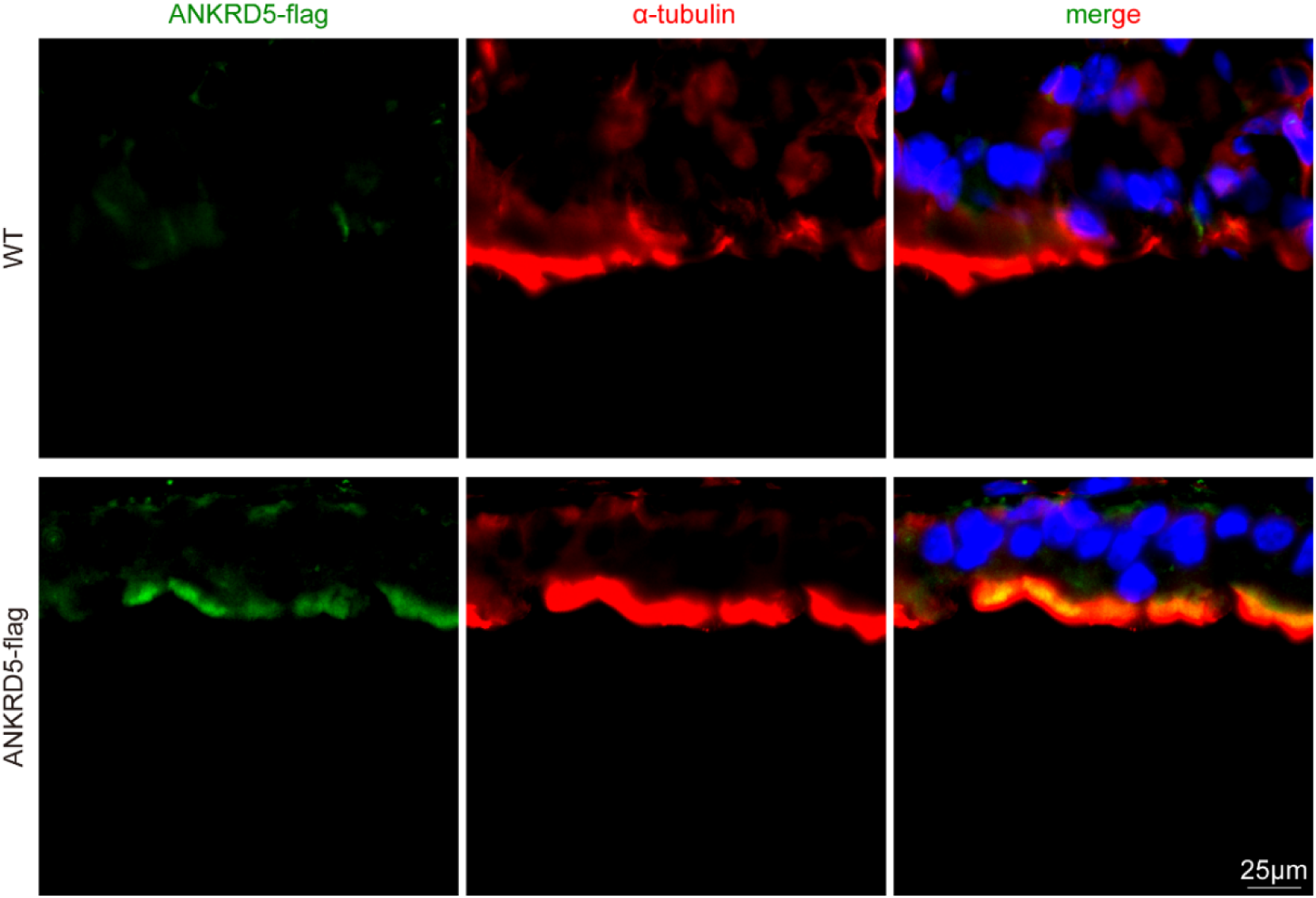
ANKRD5 was expressed in mouse respiratory cilia. Mouse respiratory cilia were double-stained with anti-α-tubulin antibody (red) and anti-Flag antibody (green), nuclei were stained with DAPI (blue). Expression of ANKRD5 protein was detected in the cilia of the respiratory tract in ANKRD5-Flag mice, and it exhibited a similar pattern to the α-tubulin signal. The scale bar represents 25μm.

**Figure S4.**
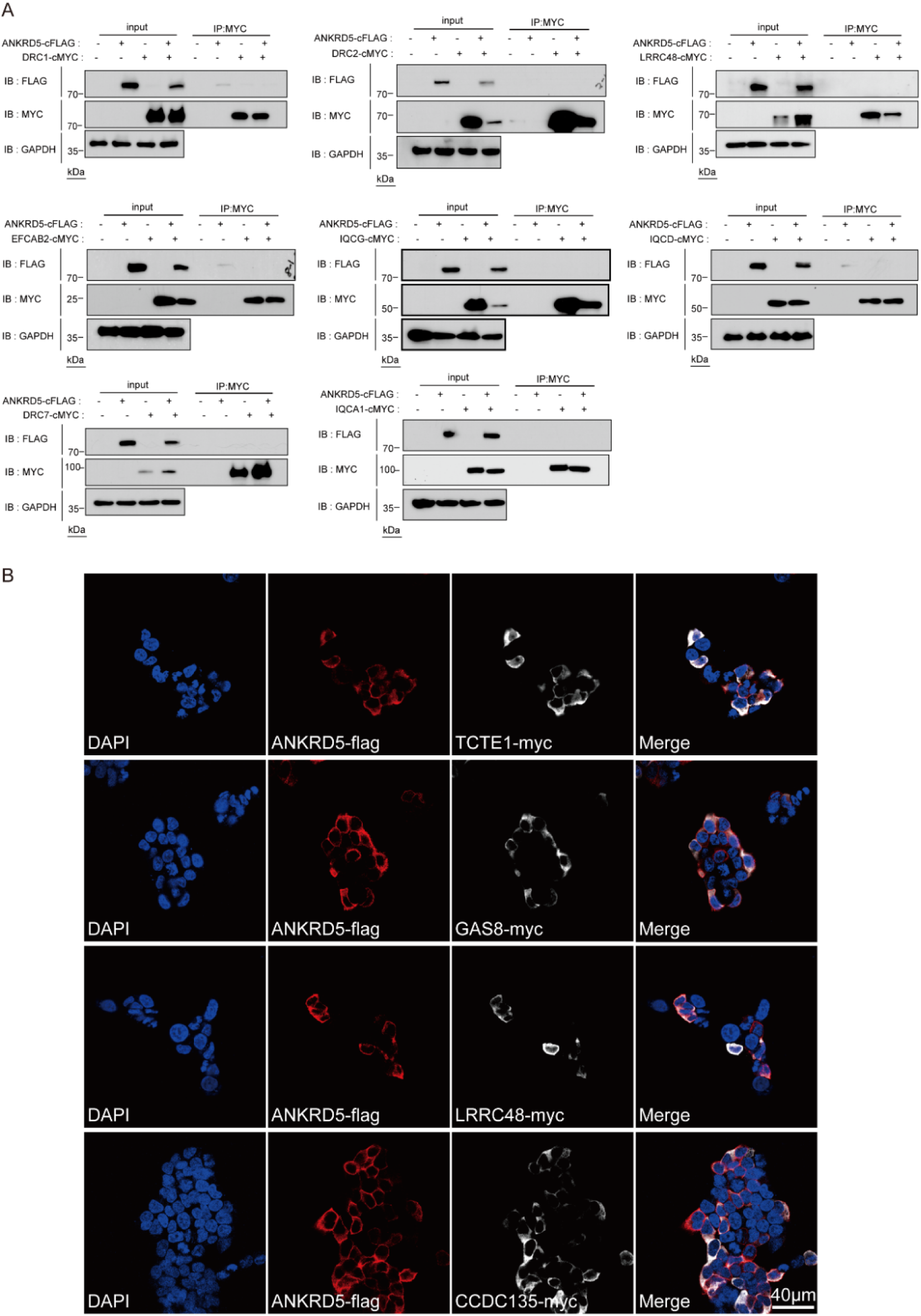
Identify the interaction of ANKRD5 and other N-DRC components. (A) Co-IP followed by WB analysis were performed using lysates collected from HEK293T cells transfected with Flag tagged ANKRD5. Immunoprecipitated proteins by anti-MYC antibody were analyzed by WB using anti-Flag antibodies. (B) HEK293T cells transiently expressing ANKRD5-Flag and DRC-MYC were stained with Flag (red) and MYC (white) to visualize ANKRD5 and DRC, respectively. DAPI (blue) was used to visualize the nuclei.

**Figure S5.**
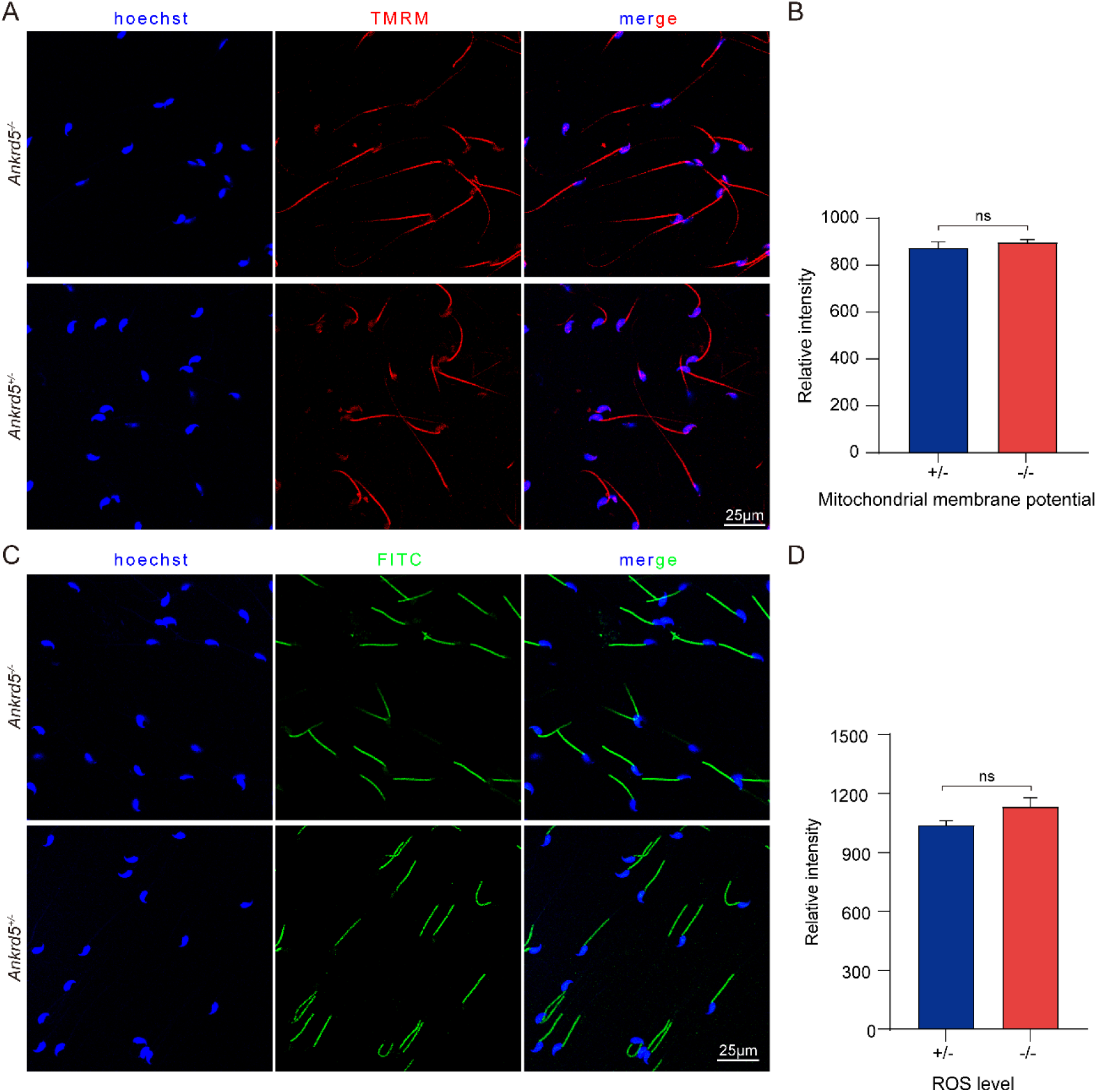
The mitochondrial membrane potential and ROS levels of *Ankrd5* null sperm was normal. (A) Mitochondrial activities assessed by fluorescence of TMRM, RFP filter. The higher the potential, the higher the concentration of TMRM in mitochondria, resulting in an increased fluorescence intensity. (B) Graphs show semi-quantitative of TMRM fluorescence intensity. Values represent mean ± SE (n=3). (C) ROS levels assessed by fluorescence of DCFH-DA, FITC filter. The higher the concentration of ROS, the higher fluorescence intensity. (D) Graphs show semi-quantitative of FITC fluorescence intensity. Values represent mean ± SE (n=3).

**Figure S6.**
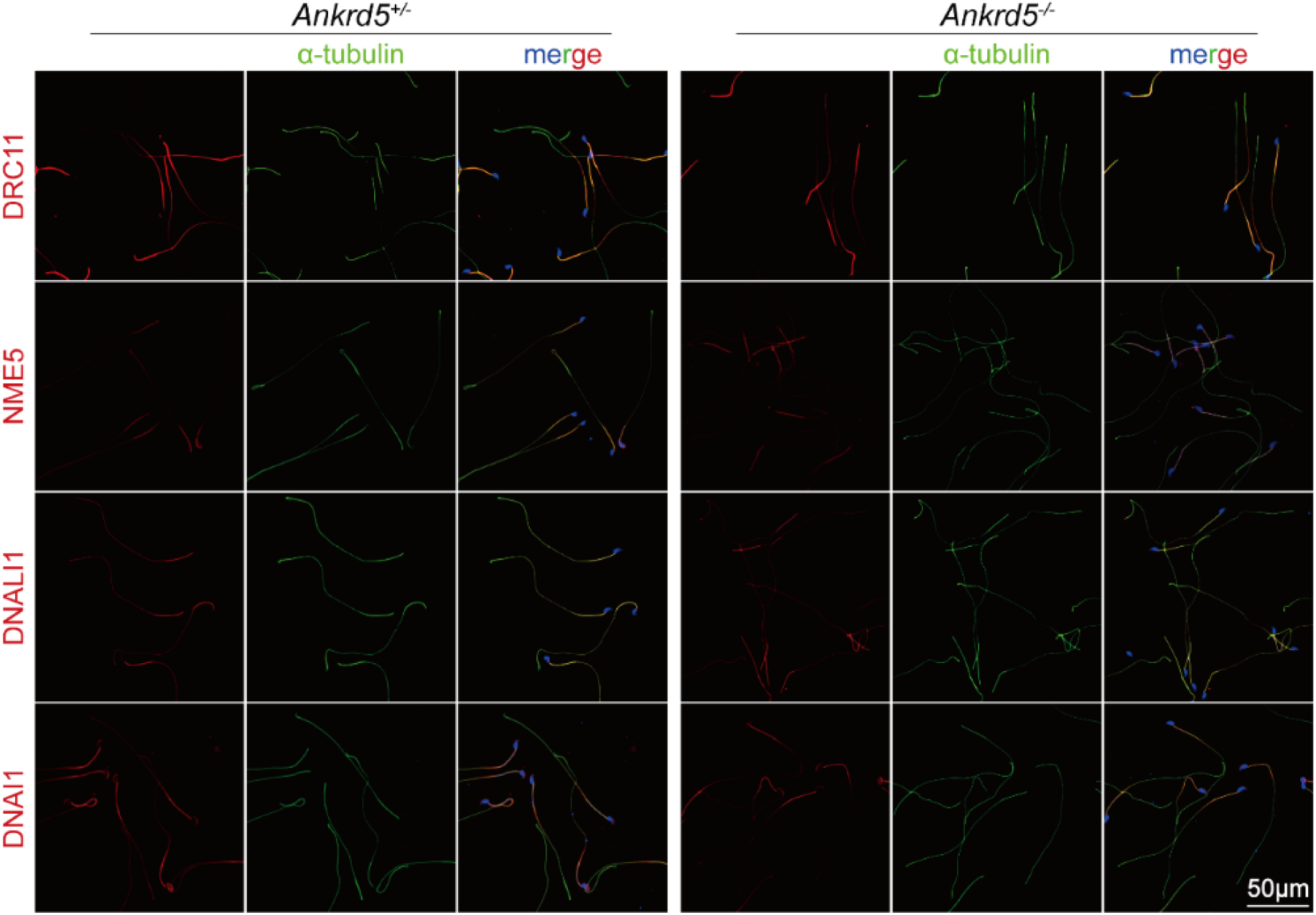
Immunofluorescence results of ANKRD5-null sperm and control. DRC11 serves as a marker protein for N-DRC (nexin-dynein regulatory complex), NME5 as a marker for RS (radial spoke), DNALI1 as a marker for IDA (inner dynein arm), and DNAI1 as a marker for ODA (outer dynein arm).

**Figure S7.**
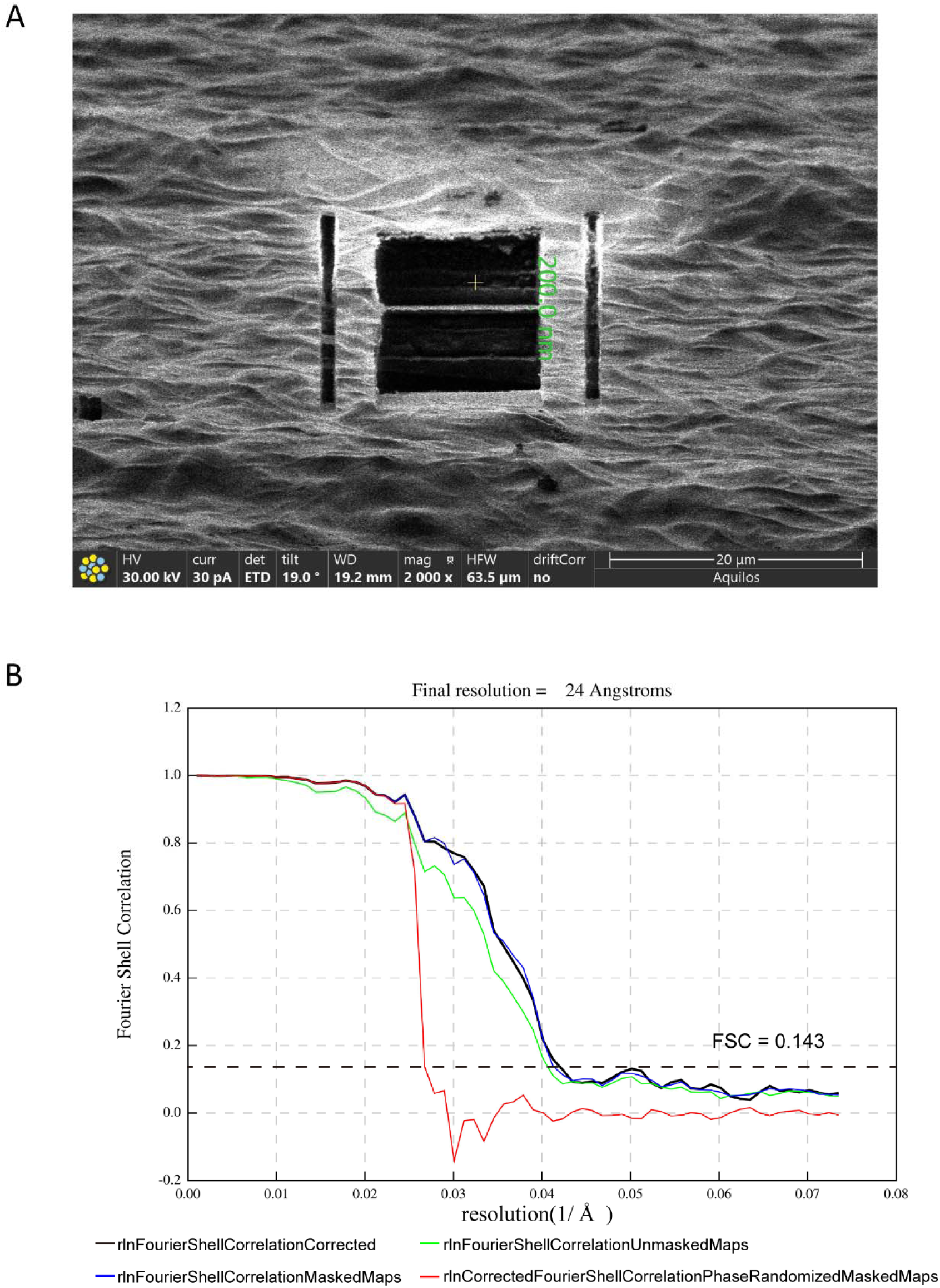
Cryo-FIB milling and the half-map Fourier Shell Correlation (FSC) of the *Ankrd5*^-/-^ mouse sperm axoneme. (A) Inspection of frozen sperm on the grid after FIB milling. The thin lamellae were used for data collection. (B) The half-map Fourier Shell Correlation (FSC) plot of *Ankrd5*^-/-^ DMT is shown.1 Å = 0.1 nm

**Figure S8.**
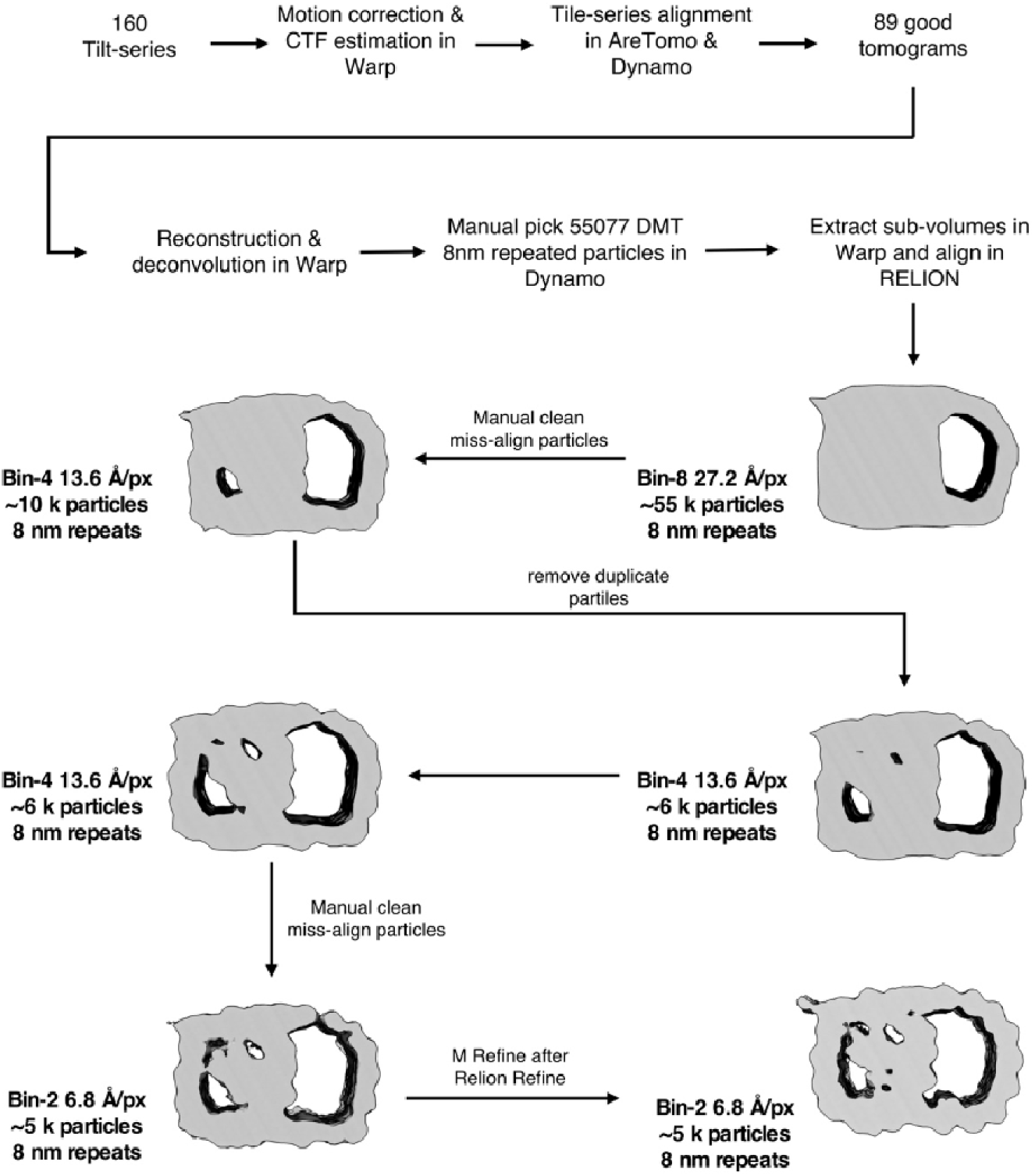
The data processing of ANKRD5-KO mouse sperm DMT. The pixel sizes at different binning levels are indicated in angstroms per pixel (Å/px for short).

**Figure S9.**
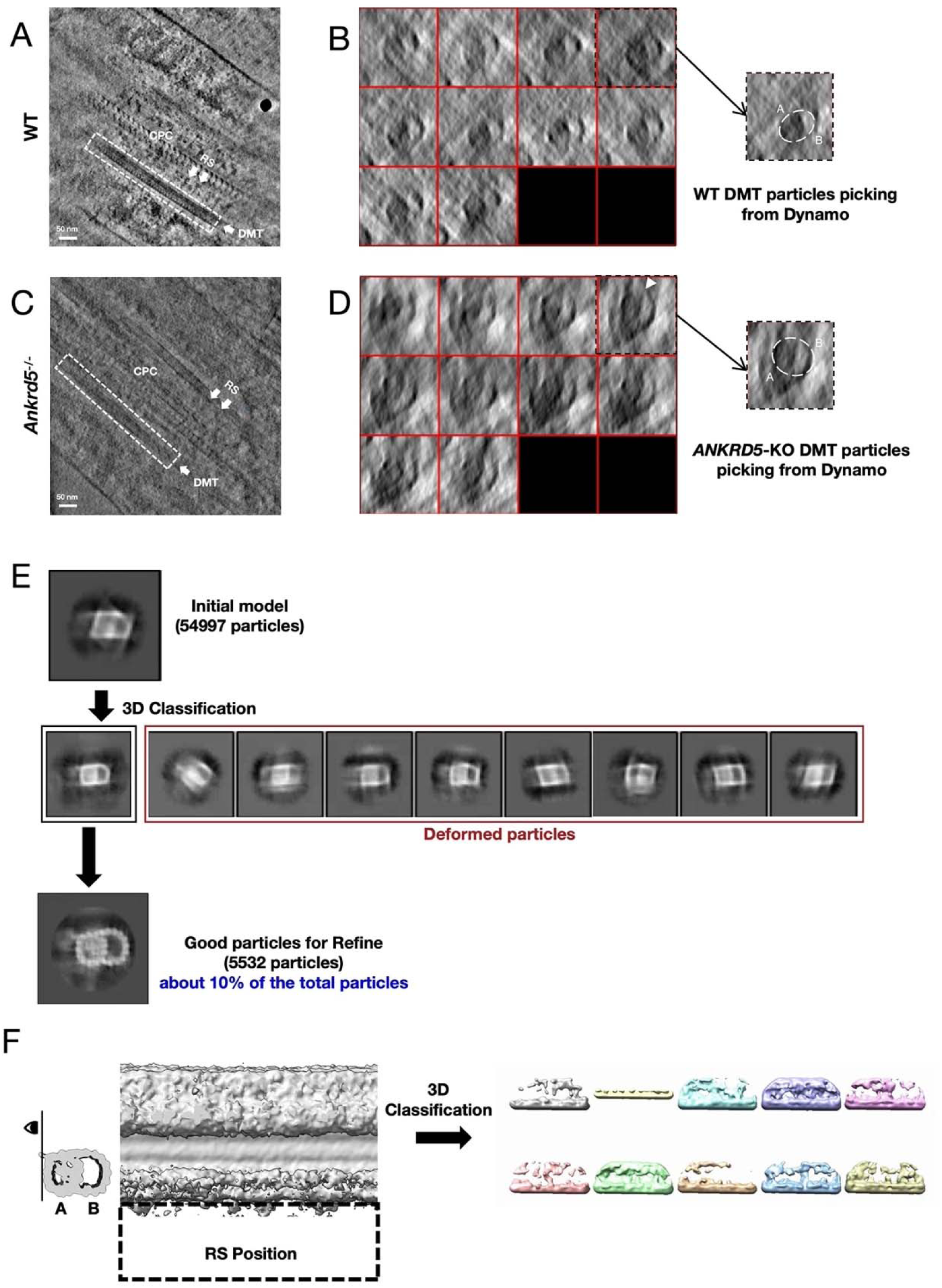
The states of DMT particles in sperm of *Ankrd5*-KO mouse. (A) and (C) Tomogram slices of WT and *Ankrd5*-KO in Dynamo (The data for WT mouse sperm was EMPIARC-200007). DMT and RS are marked with white dashed lines and white arrows, respectively. (B) and (D) Comparison of DMT particle states between WT and *Ankrd5*-KO in Dynamo. The visual angles of the DMT particles shown in (B) and (D) show that the DMT fibers within the white box in (A) and (B) are divided equally into 10 slices along the direction of the white arrow, respectively. The DMT particle shapes of WT and *Ankrd5*-KO are marked by white dashed lines on the right of (B) and (D). The white arrow in (D) identifies the junction of A-tube and B-tube that is suspected to be disconnected. (E) Deformed particles discarded in 3D classification and final aligned DMT artifacts. (F) 3D classification of attempted RS locations.

**Figure S10.**
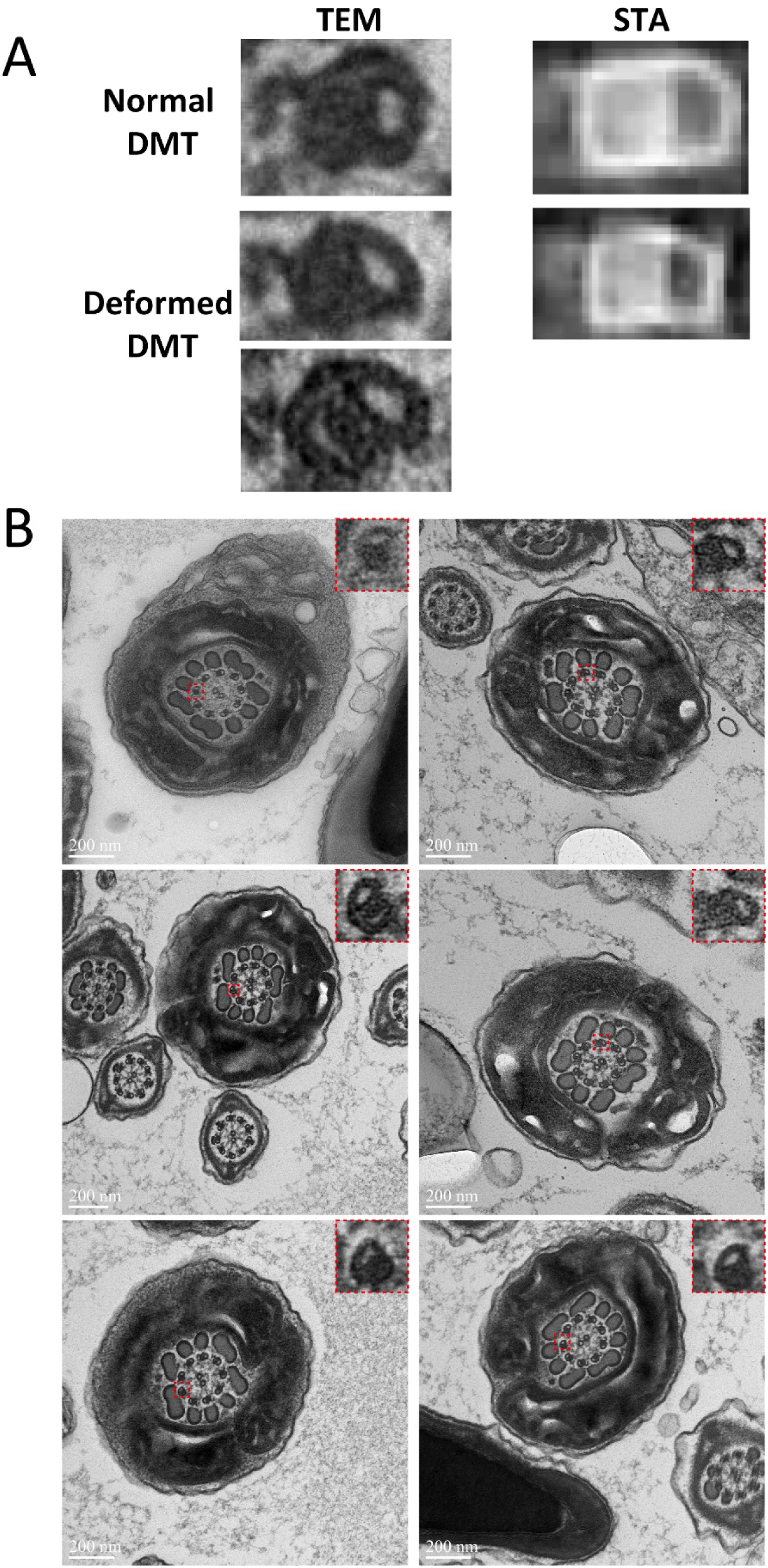
The deformed DMT in the TEM results. (A) The presence of both normal and deformed DMTs in the TEM and STA results is shown. (B) The deformed DMTs in the TEM results are all marked by red dashed boxes, and the TEM results presented in (A) are all cropped from (B).

**Movie S1.** Sperm from *Ankrd5^+/-^* mouse. Video recording of sperm from *Ankrd5^+/-^* mouse from the Computer Assisted Sperm Analyzer system (Version.12 CEROS, Hamilton Thorne Research). Capture rate was set at 60 frames/second.

**Movie S2.** Sperm from *Ankrd5^-/-^* mouse. Video recording of sperm from *Ankrd5^-/-^* mouse from the Computer Assisted Sperm Analyzer system (Version.12 CEROS, Hamilton Thorne Research). Capture rate was set at 60 frames/second.

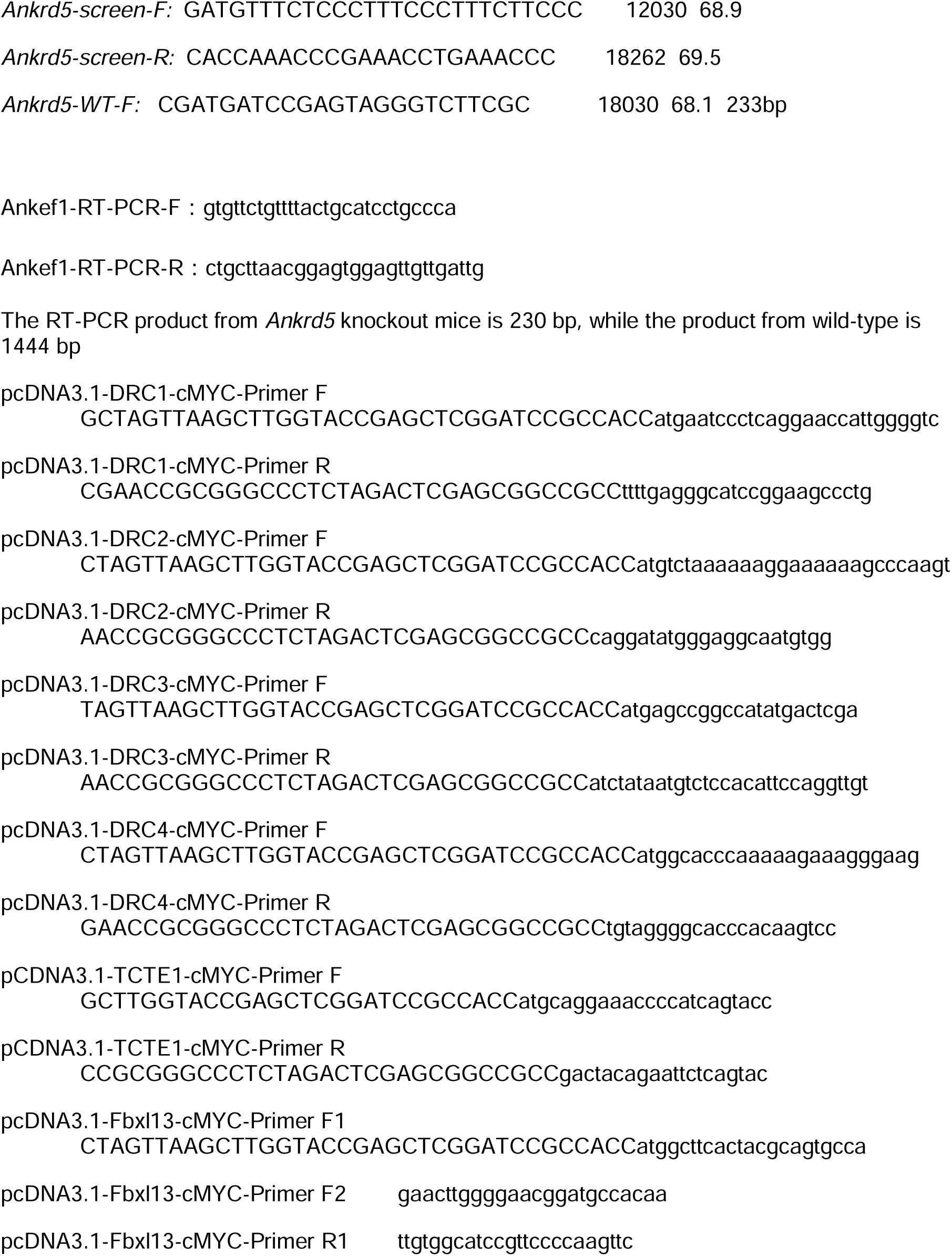
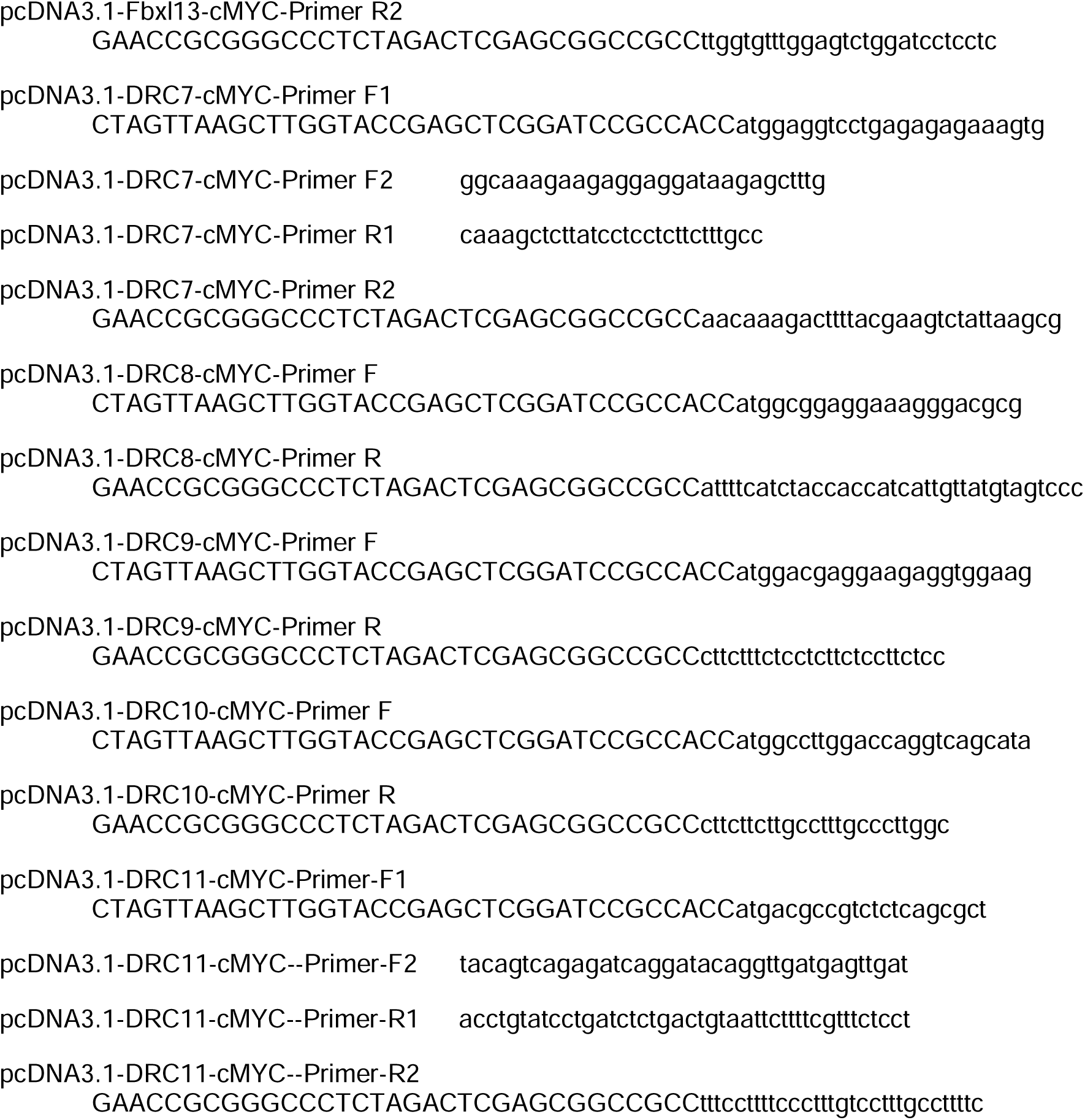

## Notes

### Competing Interest Statement

The authors have declared no competing interest.

### Summary of Updates

1. Added Figure S10. 2. Corrected the y-axis label in Figure 3G. 3. All p-values have been reformatted to italic lowercase letters to comply with the journal style guidelines. 4. Figure 6 Legend: Corrected a typographical error in the figure legend regarding the genotype comparison. 5. In the original Figure 4E, a zoom-in panel was added to highlight the deformed DMT. 6. The writing has been polished throughout the manuscript. 7. Adjusted some of the references.

